# An altered balance of integrated and segregated brain activity is a marker of cognitive deficits following sleep deprivation

**DOI:** 10.1101/2020.11.28.402305

**Authors:** Nathan E. Cross, Florence B. Pomares, Alex Nguyen, Aurore A. Perrault, Aude Jegou, Makoto Uji, Kangjoo Lee, Fatemeh Razavipour, Obaï Bin Ka’b Ali, Umit Aydin, Habib Benali, Christophe Grova, Thien Thanh Dang-Vu

## Abstract

Sleep deprivation (SD) leads to impairments in cognitive function. Here, we tested the hypothesis that cognitive changes in the sleep-deprived brain can be explained by information processing within and between large-scale cortical networks. We acquired functional magnetic resonance imaging (fMRI) scans of 20 healthy volunteers during attention and executive tasks following a regular night of sleep, a night of sleep deprivation, and a recovery nap containing non-rapid eye movement (NREM) sleep. Overall, sleep deprivation was associated with increased cortex-wide functional integration, driven by a rise of integration within cortical networks. The ratio of within vs between network integration in the cortex increased further in the recovery nap, suggesting that prolonged wakefulness drives the cortex toward a state resembling sleep. This balance of integration and segregation in the sleep-deprived state was tightly associated with deficits in cognitive performance. This was a distinct and better marker of cognitive impairment than conventional indicators of homeostatic sleep pressure, as well as the pronounced thalamo-cortical connectivity changes that occurs towards falling asleep. Importantly, restoration of the balance between segregation and integration of cortical activity was also related to performance recovery after the nap, demonstrating a bi-directional effect. These results demonstrate that intra- and inter-individual differences in cortical network integration and segregation during task performance may play a critical role in vulnerability to cognitive impairment in the sleep deprived state.

**Significance Statement:** Sleep deprivation has significant negative consequences for cognitive function. Understanding how changes in brain activity underpin changes in cognition is important not only to discover why performance declines following extended periods of wakefulness, but also for answering the fundamental question of why we require regular and recurrent sleep for optimal performance. Finding neural correlates that predict performance following sleep deprivation also has the potential to understand which individuals are particularly vulnerable to sleep deprivation, and what aspects of brain function may protect them from these negative consequences on performance. Finally, understanding how perturbations to regular (well-rested) brain functioning affect cognitive performance, will provide important insight into how underlying principles of information processing in the brain may support cognition generally.

## Introduction

The cognitive consequences of acute total sleep deprivation (SD) are substantial, negatively affecting a wide range of processes including attention, vigilance and working memory (Alhola et al. 2007, Killgore 2010). Yet the measurable impact of SD on cognition varies across individuals in a trait-like manner (Rupp et al. 2012, Dennis et al. 2017). Investigating intra-individual changes in brain activity patterns has significant potential in understanding the inter-individual effects of prolonged wakefulness on cognitive functioning. Previous studies have demonstrated the effects of acute total SD on brain activity measured with functional Magnetic Resonance Imaging (fMRI) during the resting-state, such as disrupted connectivity within and between large scale cortical networks (Yeo et al. 2015), and a reduction in network modularity (Ben Simon et al. 2017). However, very few studies have investigated the changes in brain connectivity during the performance of cognitive tasks.

Both localized brain regions and cortical networks display characteristic and independent (segregated) activity patterns during the performance of tasks (Fox et al. 2012, Di et al. 2013). However, even simple behaviors and cognitive processes require the coordination (integration) of information flows across systems distributed in the brain (Varela et al. 2001). It has been proposed that efficient cognitive functioning is reflected through a balance of segregation and integration of information across brain networks (Sporns et al. 2004, Tononi et al. 1994).

Endogenous activity fluctuations in the human brain share a robust association with ongoing cognitive and perceptual processes (Podvalny et al. 2019, He 2013), as they may interfere with the capacity for information processing. Importantly, both local and global fMRI activity fluctuates at a significantly greater magnitude in states of reduced arousal and vigilance (Wong et al. 2013), as well as following SD (Yeo et al. 2015, Nilsonne et al. 2017, Wang et al. 2016). Therefore, in states of reduced arousal, an elevation in endogenous fluctuations across the cortex may affect the ability to integrate information between and within cortical networks, corresponding to the cognitive deficits experienced during these states. These deficits may also be driven by certain changes in subcortical regions, such as the thalamus, which is involved in both maintaining cortical arousal (Van der Werf et al. 2002) and constraining the engagement of distributed neural assemblies within the cerebral cortex (Shine 2020). However, this has not been explored following experimental manipulation of arousal states, such as SD. Furthermore, SD induced cognitive impairment partially improves with subsequent sleep (Lo et al. 2016). Yet it remains unclear whether this behavioural recovery is associated with a functional ‘recovery’ of cortical networks after sleep.

Here we assessed principles of functional brain activity across various arousal states using simultaneous EEG-fMRI recording and a sleep deprivation protocol. Participants were scanned both at rest and whilst performing cognitive tasks probing attention, executive control and working memory, after both a regular night of sleep and 24hrs of SD. Additionally, we measured EEG-fMRI activity during a recovery nap containing non-rapid eye movement (NREM) sleep following SD, as well as during a post recovery nap (PRN) session when the rest and task sequences were repeated. Firstly, to quantify the functional interactions within and between cortical networks across vigilance states, we computed a measure of integration of fMRI activity (Boly et al. 2012, Marrelec et al. 2008) within and between functional networks across different hierarchical levels of a representation of the cortex. We then compared the changes in functional integration during task performance to the magnitude of cognitive impairment following SD. In addition, we measured the association between recovery of both performance and the integration of cortical networks following the nap. Next, we assessed these changes in the context of changes to the amplitude of both local activity fluctuations and the global brain signal. Finally, we investigated the influence of thalamocortical activity on functional integration in the cortex across different arousal states.

## Results

Twenty participants (mean age of 21.2 ± 2.5 years, 12 females) participated in the study. In each session (Rested Wakefulness, RW; Sleep Deprivation, SD; Post Recovery Nap, PRN) participants completed a 5-minute resting state sequence (fixation cross) (Fig. 1A). Participants also completed 3 cognitive tasks (Fig. 1B). Two were focused on attention - the Attention Network Task (ANT (Fan et al. 2002), 13min) and the Mackworth Clock Task (MCT (Lichstein et al. 2000), 5min). The other focused on working memory - the N-back task (Kirchner 1958) (8min). As expected, performance outcomes (accuracy and reaction time) across all tasks were significantly impaired following SD, and improved following the recovery nap (SI Results, Fig. S1, Table S1).

**Fig 1.**
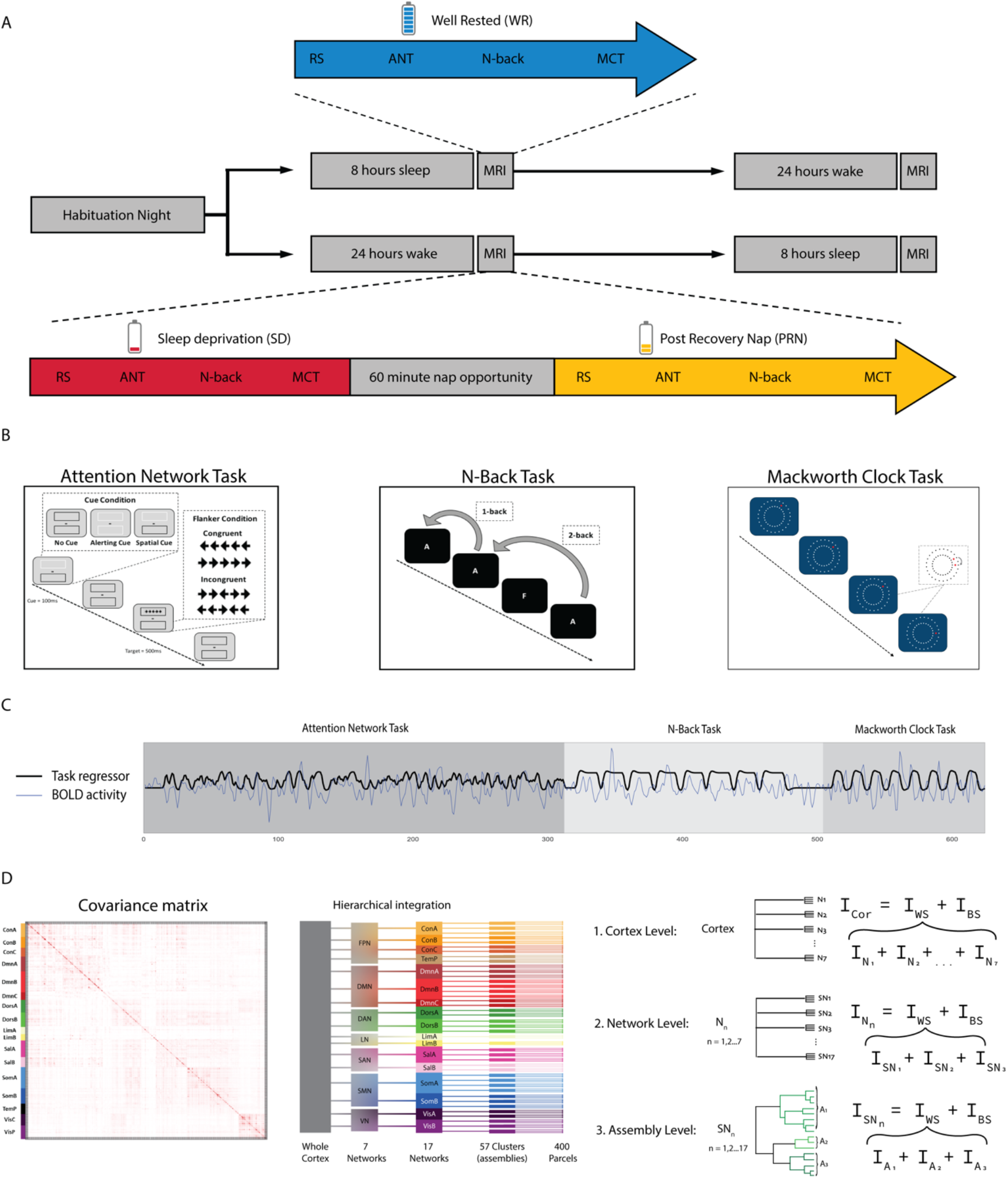
**(A)** Study design and behavioural results. Participants made 3 visits to the lab: a habituation night, followed by a counterbalanced design of either another full-night opportunity to sleep (blue), or a night of total sleep deprivation. In the morning following each night, participants completed resting state (RS) and cognitive tasks inside the MRI scanner. In the sleep deprived state (red), participants also had a recovery nap opportunity (yellow) and then recompleted the tasks inside the MRI scanner. **(B)** Participants completed three cognitive tasks inside the scanner in each condition: the Attention Network Task (ANT), N-back task and Macworth Clock Task (MCT). **(C)** The preprocessed BOLD time series was concatenated across the three tasks, providing one timeseries for each subject and condition. Task-specific activity (trials for the ANT, blocks for the N-back and MCT) was modelled with a finite impulse response combined with a hemodynamic response function (black line) and regressed from the timeseries (blue line) for each parcel to reduce the influence of task-evoked coactivation in subsequent connectivity analyses. **(D)** Integration was calculated from the covariance matrix of cortical parcels, in a hierarchical manner: across the whole cortex, 7 networks, 17 networks, and 57 clusters. At the cortex level, total integration was calculated as the sum of integration within each of Yeo7 networks, and a single measure of integration between networks (between systems). The total integration within each Yeo7 network was calculated as the sum of integration within each of Yeo17 sub-networks; and the integration within each of the Yeo17 networks as the sum of integration within each cluster of parcels, as well as a single measure of integration between sub-networks or clusters.

**Fig 2.**
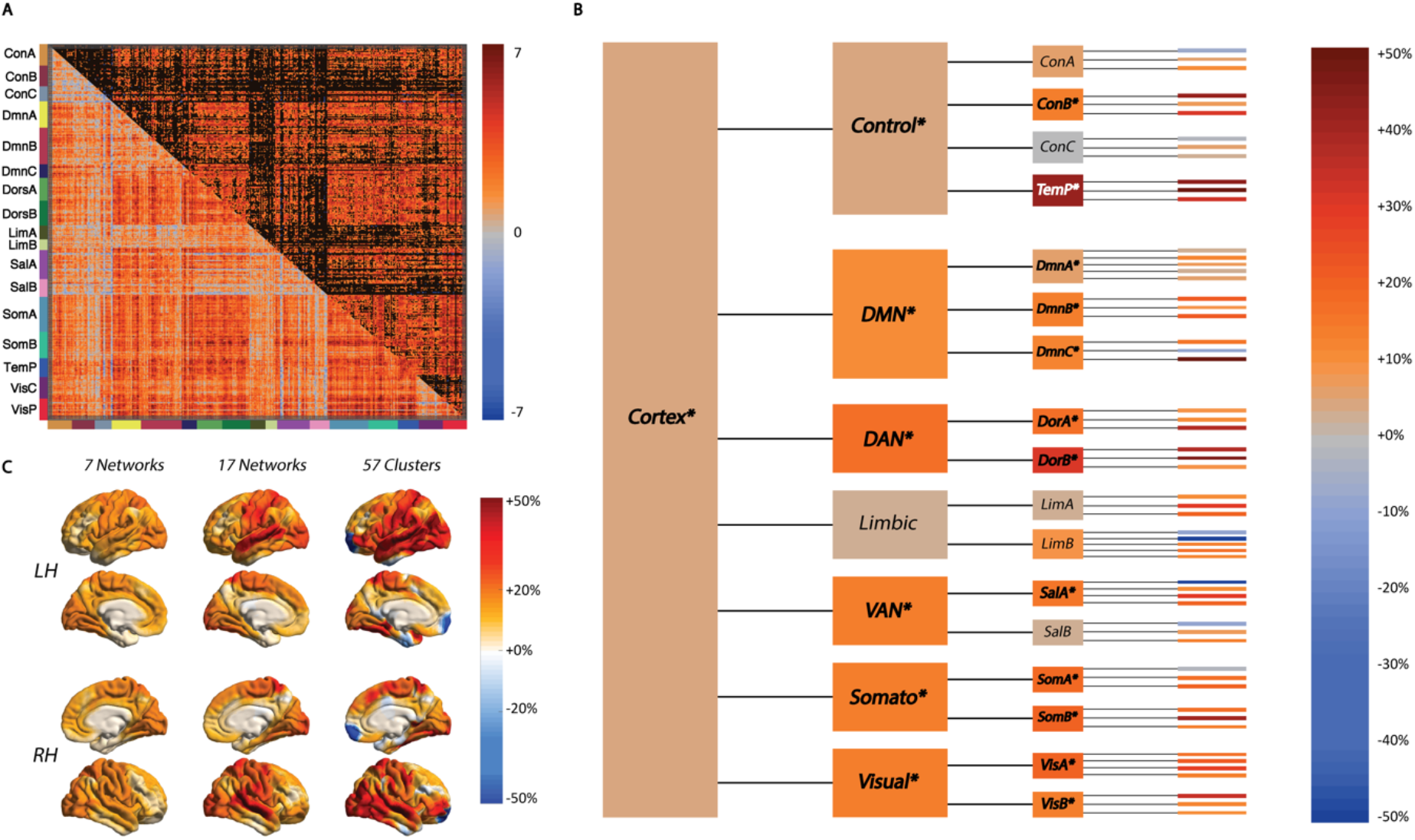
**(A)** The t-test matrix of change in functional correlations between the timeseries of 400 cortical parcels (during task performance) depicts a widespread increase in correlations between sleep deprivation (SD) and well rested (WR) states (unthresholded, lower triangle; thresholded PFDR<0.05, upper triangle). (B) Changes in integration (as a percentage of the value in the WR condition) are shown across different levels of a hierarchical model of the cortex: whole cortex, 7 networks, 17 networks, and 57 clusters. The total integration within the cortex increased from the WR to the SD state, but not within all networks (*italics depict significant changes measured through a Bayesian framework). (C) The % change in total integration at each level of the hierarchy mapped onto the cortical surface illustrates the increase in integration is most focused toward the centre of the lateral surface. DMN = Default Mode Network; DAN = Dorsal Attention Network; VAN = Ventral Attention (Salience) Network; Somato = Somatomotor Network

Through EEG confirmation, 13 participants (65%) had brief sleep episodes (<10s) and 4 (20%) fell asleep (>30s) during the resting state sequences in the SD condition. In addition to excessive head motion (average framewise displacement: 0.16 ± 0.09mm in RW vs 0.22 ± 0.14mm in SD RW), this resulted in these sequences not being used in the main analyses, however results from the resting state data can be found in (SI Results, Fig. S2). None of the subjects in the final sample fell asleep during the cognitive tasks. Additionally, head movement inside the scanner during the cognitive tasks was not significantly different between the sessions (average frame-wise displacement across 3 tasks: 0.15 ± 0.08mm (RW) vs 0.17 ± 0.09mm (SD); F = 1.06, p = 0.301). Therefore, we focused our main analyses on task fMRI data during the RW, SD and PRN states, as well as during the recovery nap consisting of NREM sleep.

The fMRI data across the multiple tasks were concatenated temporally for every subject, resulting in a 26min long time series of blood oxygen level dependent (BOLD) signal (Fig. 1C). The general task-specific activity (i.e. for all trials for the ANT and all blocks for the MCT and N-back tasks) was regressed out from the BOLD time series as has been implemented in previous studies (De Havas et al. 2012, Wang et al. 2016) to reduce the influence of task-evoked activations in connectivity and integration analyses. The BOLD data sampled along the cortical surface was clustered into 400 parcels (Schaefer et al. 2018). Integration was calculated at different levels in a constructed hierarchical model of the cortex. The whole cortex was divided into 7 networks and each of these 7 networks were further divided into smaller sub-networks (17 networks in total), based on a widely used functional template of cortical networks (Yeo et al. 2011). We further divided the 17 sub-networks into smaller clusters (57 clusters in total, estimated through hierarchical clustering within each of the 17 sub-networks, Fig. 1D, S3) to assess integration changes at an even more localised level than 17 networks. Total integration (***I***_*Tot*_) equates to the sum of within and between sub-system integration (i.e. within and between 7 networks at the whole cortex level). Differences in total integration were compared between the three vigilance states and NREM sleep within a Bayesian framework as has been used previously (Boly et al. 2012, Marrelec et al. 2008).

### Altered Functional Integration in the Cerebral Cortex during the Sleep Deprived State

Overall, there was a widespread increase in correlation values (functional connectivity) between cortical parcels from RW to SD (41% of edges significantly increased, **Fig. 1*A***). This coincided with an increase in the (***I***_*Tot*_) measured across the entire cortex. **Fig. 1*B*** illustrates the changes in functional integration across distinct levels in the hierarchical model of the cortex. At the level of 7 cortical networks, there was an increase in integration within 6 of the 7 networks, with the exception of the Limbic network (Table S2). At the 17-network level, 12 networks demonstrated increased integration, while one of the Ventral attention (B), and two of the Frontoparietal Control (A & C) and Limbic networks showed no change (Table S3). At the cluster level, there was more of a variety of change in integration, and two clusters within the Limbic network demonstrated a substantial decreased integration from RW to SD. This pattern of change indicates that while increases in total integration were detectable when measured across the entire cortex, this was not uniform and preferentially affected certain networks and regions at lower, more localised, levels of the hierarchy (**Fig. 1*C***).

Further deconstructing these changes in integration following SD, we additionally calculated the degree of functional clustering within the cortex. Functional clustering estimates how integration is hierarchically organized within and between the constituent parts of a system, such as networks in the cortex. Specifically, the functional clustering ratio (FCR) is the ratio of within-network integration to between-network integration, and quantifies the degree of functional segregation of a given system into subsystems (Boly et al. 2012). There was a significant increase in the FCR from RW to SD at both the level of the entire cortex, and within 4 of the 7 cortical networks: Control, Default Mode, Somatomotor and Ventral Attention (Table S4). The increase in both ***I***_*Tot*_ and FCR were highly correlated across subjects (r=0.92, p<0.001). This indicates that the increase in ***I***_*Tot*_ following SD was driven by integration within each network than any increased integration between networks (Figure S4). After parsing each of the 17 networks into assemblies (clusters) based upon a hierarchical clustering of each network (Fig. S3), the only significant increases in the FCR were observable within the Control (A), Default Mode (C), Somatomotor (B) and Ventral Attention (A) networks (Table S5).

To demonstrate the robustness of these findings to the choice of preprocessing or parcellation, we replicated these analyses without regressing out the task-specific activity (Table S6), or when using network communities defined by a data-driven clustering of the functional connectome (via the Louvain algorithm(Rubinov et al. 2011)) rather than an a priori enforced template (Table S7, Fig S5). These analyses support the main findings. However, more cortical networks demonstrated significant increases in integration induced by SD when the task evoked activity were not regressed out (7 of the Yeo7 networks, and 15 of the Yeo17 networks), suggesting that task activations may interact with sleep deprivation to further drive an increase in cortical within-network integration.

### Altered Functional Integration in the Cerebral Cortex is Related to Changes in Cognitive Performance

To assess how the changes in brain activity patterns related to changes in cognitive performance, we first assessed a combination of performance change across all tasks. The difference in performance outcomes (SD-RW) on each of the 3 separate cognitive tasks were grouped into either accuracy (% of correct responses) or speed (reaction time, ms). These change scores for accuracy or speed were entered into two separate Principal Component Analyses (PCA), to obtain resulting the principal components that explained the most variance in Accuracy or Speed across the 3 tasks. The first component for Accuracy (explaining 71% of the variance across tasks) was robustly and significantly negatively associated with the change in FCR within the cortex (**Fig. 3*A***). The first component for Speed (explaining 95% of the variance) was also significantly positively correlated with the change in FCR across the entire cortex. Very similar results were observed for associations between the increase in (***I***_*Tot*_) and decrease in performance (SI Results, Fig. S6). The change in integration within each of the 7 networks was negatively correlated with change in Accuracy and Speed from RW to SD (SI Results). We also assessed the change in performance on each task separately in comparison to the change in (***I***_*Tot*_) during that task only. A greater increase in (***I***_*Tot*_) was significantly related to worse accuracy and speed in the ANT and Nback tasks, but not the MCT (Fig. S7).

**Fig 3.**
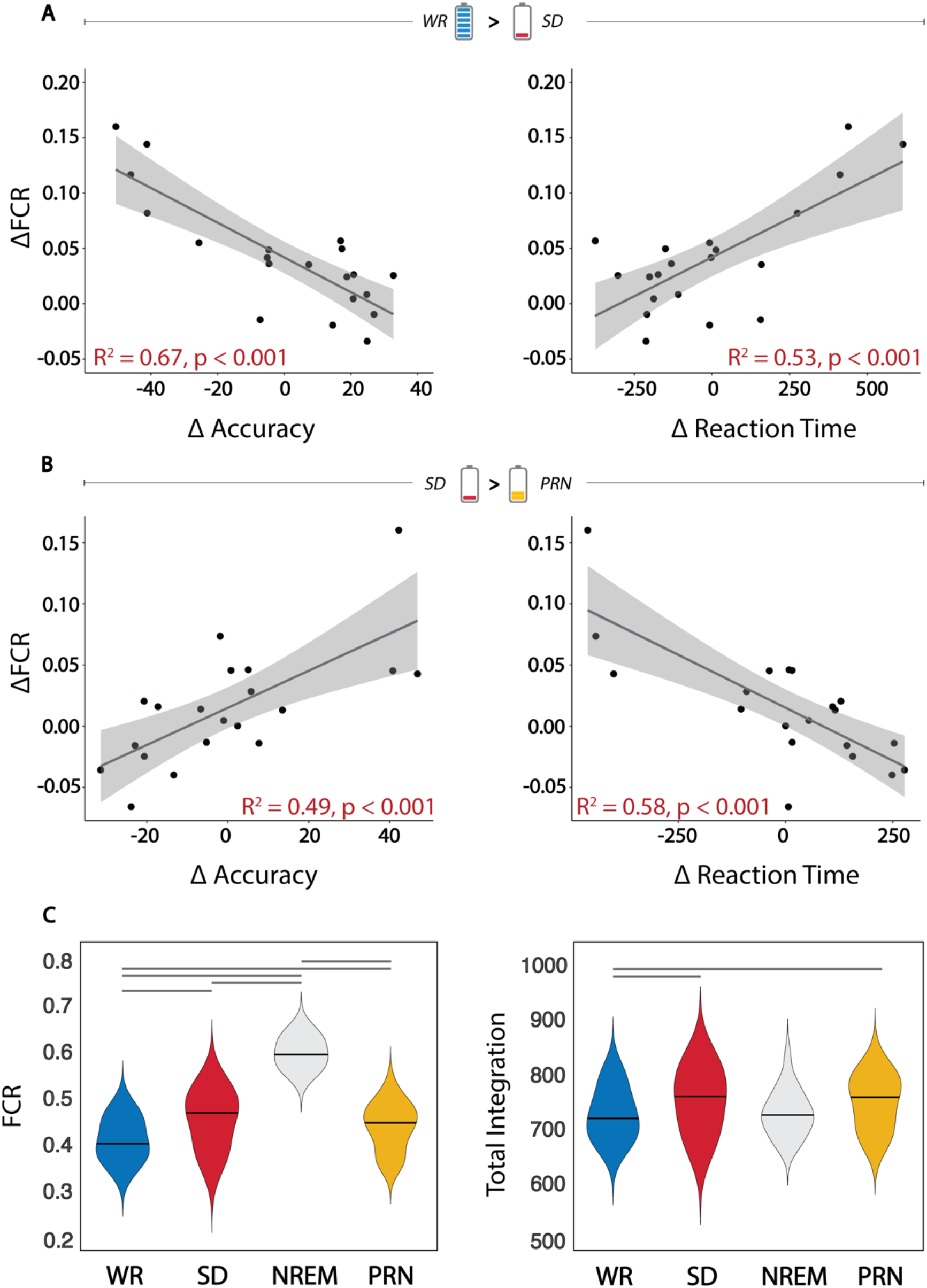
**(A)** There is a strong relationship between the change in total integration across the cortex and decrement in both performance accuracy and speed across three cognitive tasks from the WR to the SD state (top panel; performance on the x-axis is depicted as the first component from a principal component analysis on task outcomes). **(B)** The paired change in integration and performance was also observed for recovery after the nap. **(C)** The functional clustering ratio (FCR) increased from RW to SD, however significantly increased in all subjects during the nap containing NREM sleep, indicating the role of FCR in conscious states (lines represent significant within-subjects differences between sessions) **(D)** Conversely, total integration increased within-subjects from RW to SD, but there was a drop during the recovery nap despite remaining high upon waking after the nap (lines represent significant within-subjects differences between sessions).

### Changes in functional Integration and Cognitive Performance Following a Recovery Nap

A decrease in connectivity was observed from the SD to the PRN state, although to a lesser extent than the increase observed between RW and SD (16.7% of connections significantly decreased, Fig. S9). There was a small decrease in FCR across the entire cortex (**Fig. 3*B***). This was not correlated with the duration of the nap or the amount of slow wave sleep across individuals (SI Results). However, the subjects who had the greatest decrease in FCR from SD to the PRN, showed the largest improvement in performance for both the Accuracy and Speed outcomes (**Fig. 3*A***). This recovery effect demonstrates a bi-directional link between changes in functional integration and changes in cognitive performance due to SD.

### Functional Clustering of Brain Activity Is Robustly Associated with Conscious States

Following the SD condition, 19 of the 20 participants managed to successfully sleep inside the scanner, for an average of 50 ± 12 minutes (a sleep summary can be found in Table S8). To ensure reliability of integration measures in all conditions (RW, SD, PRN and sleep), a stable period of 26 minutes (i.e. the same length as the concatenated task time series) consisting of NREM sleep (Stages N2 and N3) was selected from the fMRI scan sequence, and the same connectivity and integration metrics were calculated. During this recovery nap of NREM sleep, the FCR was significantly greater during NREM sleep than the wake states, for all subjects (**Fig. 3B**). This was despite the amount of (***I***_*Tot*_) being comparable to the other wake states (**Fig. 3C**). The increase in FCR and stability of (***I***_*Tot*_) was driven by both an increase in within-networks integration as well as a decrease in the integration between-networks during the nap (SI Results, Fig S4). These changes demonstrate that the altered balance of within-networks and between-networks integration is a characteristic feature of online vs offline brain states.

### Functional Integration is Linked to the Amplitude of Local and Global Signal Fluctuations

We investigated whether the increase in functional integration may be due to regional changes in the amplitude of brain activity fluctuations. Between the RW and SD states, the parcel-wise change in signal fluctuation amplitude (standard deviation of task-regressed signal in each 400 parcels) was greatest over the somatosensory cortex and peripheral visual cortex (**Fig. 4*A***). Across individuals these changes in signal fluctuation amplitude were only correlated with individual-level changes in integration within the visual cortex (**Fig. 3*B***). However, the spatial correspondence between the group-level changes in signal fluctuation and changes in integration at the group level were all significantly correlated, and the relationship was strongest for integration changes at the cluster (assembly) level (**Fig. 3*C***).

**Fig 4.**
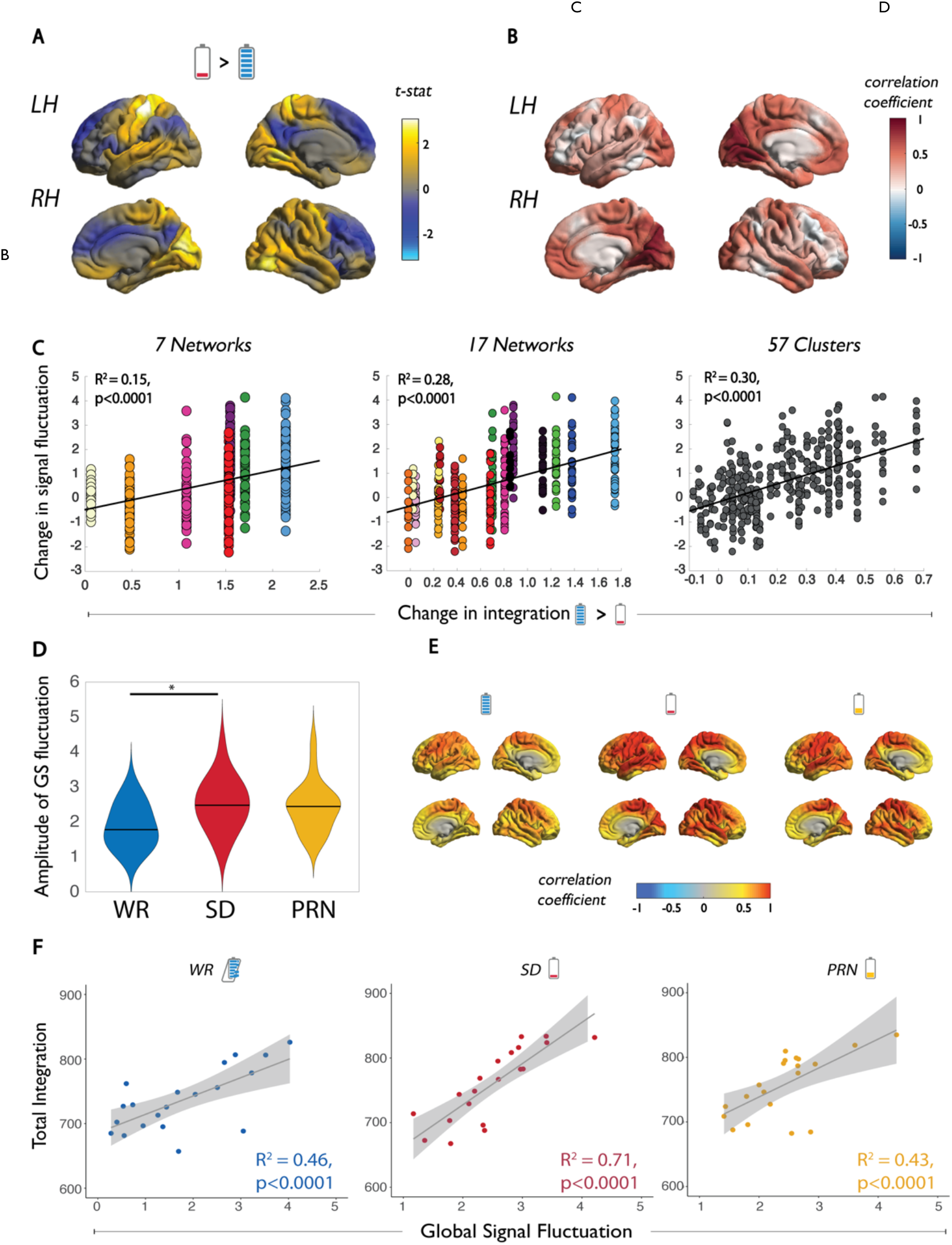
The relationship between functional integration and local and global signal fluctuations. **(A)** The group-average change in amplitude of signal fluctuation (standard deviation of signal) in each of 400 cortical parcels between SD and WR states. **(B)** The individual-level correlations between change in amplitude of the signal fluctuation and integration within 17 networks. **(C)** Correspondence between the group-average change in amplitude of signal fluctuation with group-average total integration within 7 networks, 17 networks, and 57 clusters (significance corrected with 100 spin permutations). **(D)** The distribution across subjects of the amplitude of the global signal fluctuation in the WR, SD and PRN states. **(E)** Group-average correlation coefficient between each parcel’s time series and the global signal in each state. **(F)** Individual relationships between total integration across the cortex and the amplitude of the global signal fluctuation in the WR, SD, and PRN states.

In line with previous reports (Nilsonne et al. 2017, Wang et al. 2016, Yeo et al. 2015), the amplitude of the global signal fluctuation (standard deviation of the global signal time series) significantly increased from the RW to the SD state (**Fig. 3*D***). At the RW state, the time series of each parcel were only moderately associated with the global signal time series, however this increased significantly across a widespread number of parcels, including somatomotor, visual, temporal and parietal regions of the cortex during the SD state, and persisted in the PRN state (**Fig. 3E**). In each state, the (***I***_*Tot*_) was significantly correlated with the global signal fluctuation, however the relationship was the strongest during the SD state (**Fig. 3F**). This suggests that the global signal is strongly associated with measures of functional integration, but these measures become more tightly coupled in periods of reduced arousal, such as following SD. To ensure these results were not driven by motion, the relationship between the global signal fluctuation and (***I***_*Tot*_) remained significant when average framewise displacement in each state was included as a covariate in the regression model (SI Results).

### Relationships with Sleepiness and Physiological Markers of Sleep Pressure

Given the increase in both (***I***_*Tot*_) and the FCR within the cortex following SD, we tested whether these were related to the level of homeostatic sleep pressure. Firstly, the change in (***I***_*Tot*_) was not associated with subjective sleepiness (Karolinska Sleepiness Score (Shahid et al. 2011)). Neither did the magnitude of the increase in (***I***_*Tot*_) or FCR following SD correlate with sleep latency, latency to Stage 3 NREM sleep or the amount of slow wave activity (SWA) during the recovery nap (SI Results). Additionally, despite a slight slowing of EEG frequencies from RW to SD (1-4Hz, SI Results), these changes were not related to the change in FCR or (***I***_*Tot*_). These all have been previously shown to be markers of homeostatic sleep pressure, suggesting that functional integration in the cortex is not merely a marker of enhanced sleep pressure. Furthermore, none of the aforementioned measures of homeostatic sleep pressure were related to the change in accuracy or reaction time from RW to SD (SI Results).

### Changes in Subcortical Connectivity

We also measured subcortical-cortical connectivity to investigate whether these interactions may influence changes in cortical integration in periods of reduced arousal. Out of five subcortical regions (amygdala, caudate, pallidum, putamen, thalamus) and the hippocampus, there was a notable significant decrease in the correlations between the thalamus and widespread cortical regions following SD (**Fig. 5*A***). Connectivity changes with the thalamus were greatest in the Somatomotor, Dorsal Attention and Temporal Parietal networks (**Fig. 5*B***). However, no correlations between the change in thalamocortical connectivity and cortical integration were observed across individuals (**Fig. 5*C***). Neither were there any relationships observed between thalamocortical connectivity and cognitive performance, or markers of sleep pressure (SI Results). Nevertheless, thalamocortical connectivity was lowest during the SD state, even less than during NREM sleep (**Fig. 5*D***). This was particularly accentuated in the resting state sequence following SD, a period of sleep-wake instability when the majority of subjects had sleep intrusions (**Fig. 5*D***).

**Figure 5.**
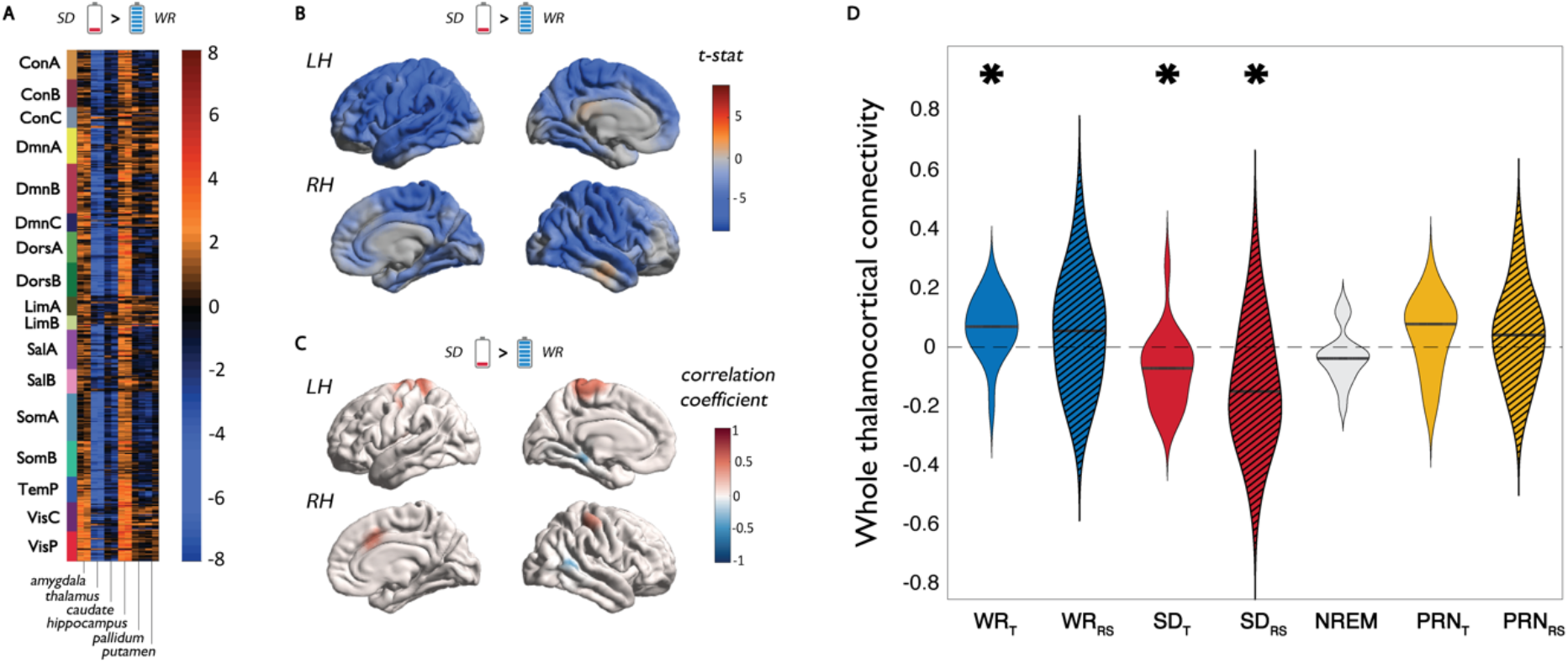
Changes in thalamocortical connectivity and the relationship to functional integration. **(A)** The group-average change in connectivity (correlation of time series) between all 400 cortical parcels and 6 subcortical regions from sleep deprivation (SD) and well rested (WR) states during task performance. **(B)** Spatial map of changes in thalamocortical connectivity between SD and WR states during task performance. **(C)** The individual level relationship between change in thalamocortical connectivity and integration within 17 networks during task performance. **(D)** The individual distributions of thalamocortical connectivity (averaged across the entire cortex) in the WR, SD and PRN states, T=Task condition; RS = Resting-state condition. *Denotes average connectivity is significantly different from zero.

## Discussion

These findings demonstrate that in states of low arousal (i.e. sleep deprivation) the integration within cortical networks increases relative to integration between networks. These changes were observed during the performance of cognitive tasks, and importantly were strongly related to overall performance. The balance of these two integration terms, i.e. the functional clustering ratio (FCR), can be viewed as a measure of the degree of functional segregation of a system into its constituent parts (Boly et al. 2012, Tononi et al. 1994). In other words, this signifies how much information cortical networks generate independently, compared with the information generated by the cortex as a whole. The FCR was found to increase across the cortex in the SD state, and preferentially within certain functional networks (each divided in turn into smaller networks; **Fig 2*A***), suggesting that these modifications in information integration were present when measured across different hierarchical levels of the cortex.

We detected the greatest FCR during the NREM recovery nap (**Fig 3*C***), consistent with previous observations during NREM sleep (Boly et al. 2012, Del Felice et al. 2015, Tagliazucchi et al. 2013). Although previous comparisons of NREM sleep were with the wakeful resting state, the current findings demonstrate that these effects generalise to other states of wakefulness (i.e. task performance). The increase in FCR was driven by both an increase in within-network integration as well as a significant decrease in between-network integration. It is widely accepted that sleep is not merely a state of quiescence, but an active brain state of self-organised, endogenous activity mostly cut-off from the outside world. The level of consciousness has been hypothesised to be related to the degree of integrated information in the brain: that is, the information generated by the interactions in a whole system, beyond the information generated by the parts (Tononi 2004, Tononi 2008). This is consistent with our observations of segregated brain activity during the recovery nap. Alternatively, in the SD state, the integration between-networks remained the same as the RW state, which is coherent with the fact that subjects remained in a conscious, wakeful state. However, given the increase of within-network integration, the FCR was still significantly higher when compared to RW. In summary, these results suggest that following prolonged wakefulness the proportional integration and segregation of brain activity within structured cortical networks appears to be driven from levels in well-rested wakefulness toward those observed during offline states, such as NREM sleep, suggesting there exists an underlying continuum of functional segregation from RW to NREM sleep. We were not able to robustly measure the differences between NREM2 and NREM3 in this study with the same amount of consecutive data points as in the wake conditions, however other studies suggest that brain networks are even more segregated during NREM3 compared to NREM2 sleep (Tarun et al. 2021), and that total integration decreases in proportion to SWA (Boly et al. 2012).

We have recently shown that the primary axes of differentiation in functional connectivity (i.e. functional gradients) across the cortex do not undergo major changes following SD (Cross et al. 2021). This is consistent with the current findings, as although within-network integration increased, between-network functional connectivity remained stable (Fig S4). Furthermore, the gradient approach is mainly sensitive to the shape of the functional connectivity distributions rather than variations in amplitude. Indeed, in this study we found that changes in the integration of brain activity are related to elevated amplitude of brain signal fluctuations, particularly the global signal (**Fig 4F**). We were able to confirm that this relationship was not due to increased motion during the SD state (SI Results). The reason behind an increase in signal amplitude during SD is not clear. However, one emerging candidate for a primary role of sleep is the flushing of metabolic waste through the mixing of cerebrospinal and interstitial fluid via the glymphatic system (Iliff et al. 2012, Xie et al. 2013). Large oscillations of fluid inflow into the perivascular space occur during sleep and are tightly coupled with large amplitude EEG slow waves (Fultz et al. 2019, Hablitz et al. 2019). Thus, a build-up of waste products from extended periods of cellular activity in the brain may trigger the mechanisms (i.e. large amplitude slow waves) for the glymphatic system to begin clearing waste, even in the wake state (Xie et al. 2013). Additionally, it is also plausible that a change in respiration or cardiac activity due to parasympathetic drive could account for changes in BOLD activity amplitude, as cerebral blood flow increases after SD (Elvsåshagen et al. 2019), and the greatest changes in integration were detected nearest to the middle cerebral artery (Gibo et al. 1981). Finally, given the assumptions of our model of integration, in particular that the BOLD data at each time point are temporally independent and identically distributed (i.i.d) realizations of a 400-dimensional random variable y (Marrelec et al, 2008), it is possible that potential sample dependencies across time could contribute to an overestimation of integration measures within the data. This could mean that the increased integration we observed in the SD condition are due to an increase in temporal dependencies in the data. This should be investigated in future studies investigating the dynamics of brain activity in the SD state.

Our results demonstrate that disruption to the balance between integration and segregation of cortical networks has a significant impact on effective and efficient cognitive performance (**Fig 3*A***), which was shown to be specific to executive attention and working memory rather than vigilance (Fig S7). This effect was bidirectional, as both increased FCR following SD negatively correlated with performance change, while the decrease in FCR following a recovery nap was associated with the greatest cognitive improvement. Maintaining a balance between integration and segregation of information flow is thought to be crucial for distributed brain networks to execute effective cognitive function (Fox et al. 2012, Tononi et al. 1994). In particular, the ability to dynamically fluctuate between integrated and segregated brain states may be the primary mechanism that supports ongoing cognitive processes (Shine et al. 2016, Sporns 2013). Information generated by the brain would theoretically decrease if dynamic states become more homogenous (indicated by an increase in total & within-network integration or global signal amplitude). Evidence from resting state fMRI studies have demonstrated that SD results in increased time spent in dynamic states associated with low vigilance (Li et al. 2020, Teng et al. 2019, Xu et al. 2018), however how SD impacts state dynamics during the performance of cognitive tasks remains to be seen.

We also replicated previous findings demonstrating that SD results in disruptive changes of thalamocortical functional connectivity, including the introduction of negative correlations (Shao et al. 2013, Yeo et al. 2015). The underlying mechanisms and physiological meaning of negative functional connectivity are still unclear (Chen et al. 2011, Fox et al. 2005, Murphy et al. 2009). However, the disruption of thalamocortical connectivity would have a significant impact on the maintenance of cortical arousal. The intralaminar and midline nuclei of the thalamus, with strong inputs from brainstem nuclei, are considered as part of the ascending reticular activating system which stimulate the cortex and facilitate the generation of intrinsic functional modes (Edlow et al. 2012, Kinomura et al. 1996, Van der Werf et al. 2002). The thalamus also serves as the major hub for gating the flow of information into the cerebral cortex by blocking incoming signals through synaptic inhibition of thalamocortical relays. This is proposed to be the main mechanism contributing to shifting the brain from an aroused state into an ‘offline’ state such as sleep (Steriade et al. 1993).

However, we observed no relationship between the change in thalamocortical connectivity and cortical integration or cognitive performance across subjects (**Fig. 5*C***). Specifically, almost every subject expressed decreased (or anti-correlated) thalamocortical connectivity in the SD state, while there were more varied changes in integration between the RW and SD states across subjects (some not exhibiting relative increases in integration or FCR after 24 hours of SD). There are two possible explanations for these findings. Firstly, during prolonged wakefulness thalamocortical disconnection may precede changes in cortical integration, the latter occurring after different durations of SD across individuals. This is supported by evidence that during sleep onset, thalamic deactivation (i.e. thalamic activity decreasing to overall sleep levels) precedes cortical changes by several minutes (Magnin et al. 2010, Spoormaker et al. 2010). This would also be consistent with the observation that the thalamocortical connectivity was at its lowest in the SD resting-state sequence (**Fig. 5*D***), when it was confirmed through EEG that the majority of subjects experienced brief entry into sleep. An alternative possibility is that disrupted thalamocortical connectivity is a state-based effect of SD, while altered cortical integration underlies inter-individual differences in cognitive performance (or cortical ‘reserves’) following SD. To compare these hypotheses would require repeated measurements of brain activity across an extended period of prolonged wakefulness. Regardless, these results further elucidate the relationship between the thalamus and cortex, demonstrating that in some individuals the cortex can still maintain an optimal balance of integration and segregation required for cognitive processing despite becoming functionally disconnected from the thalamus. As has been proposed, the ascending modulatory transmitter systems mostly provide the necessary arousal to tune the state and excitability of the different parts of the cortex, which allow for the appropriate analysis of sensory information (i.e. cognitive processing and behavioral responses; (Steriade et al. 1993). Therefore, thalamocortical disruption following SD may not directly cause impaired cognition, but rather be a precipitating factor via altered functional integration in the cortex.

## Conclusion

In conclusion, sleep deprivation appears to impact the balance of integration and segregation of brain activity. Whether these changes are completely neurogenic, or arise from systemic processes involved in maintaining homeostasis of the cellular environment in the brain remains to be elucidated. However, they appear independent from conventional markers of homeostatic sleep pressure and changes in thalamocortical connectivity. Regardless, the disruption of information integration in the brain is significantly linked to the extent of cognitive impairment experienced by individuals following SD. Future perspectives should focus on why integration and segregation of brain activity is impaired following sleep deprivation, and how this impacts the dynamics of integrated network states during cognitive task performance.

## Materials and Methods

### Population and Experimental Design

Twenty participants were recruited using advertisements posted online and within Concordia University, Montreal. All participants were 18-30 years old, healthy, regular sleepers and none were taking any medication. Participants made three visits to the sleep laboratory, including a night for habituation and screening to rule out sleep disorders, a normal night (8hr sleep, RW) and experimental night (0hr sleep, SD). Each morning in the RW and SD conditions, participants completed one resting state sequence and three cognitive tasks: the Attention Network Task, Mackworth Clock Task, and the N-back task. In the SD condition, participants were provided a 60-min opportunity to sleep inside the MRI, and the tasks and then the resting state sequence were repeated. The order of the RW and SD sessions were counterbalanced across subjects (SI Materials and Methods).

### Analysis

The change in performance outcomes on each of the 3 separate cognitive tasks were grouped into either accuracy (% of correct responses) or speed (reaction time, in ms). This provided 2 data matrices of size #subjects x #tasks; one matrix containing accuracy scores and one containing reaction times. The change scores for accuracy or speed were entered into separate Principal Component Analyses (PCA), in order to provide a global estimate of performance. The first principal component from each PCA was extracted and used as a single measure for Accuracy or Speed across the 3 tasks.

### fMRI Data Acquisition and Analysis

MRI scanning was acquired with a 3T GE scanner (General Electric Medical Systems, Wisconsin, US) using an 8-channel head coil. Structural T_1_-weighted images with a 3D BRAVO sequence, and functional echo planar images were acquired. Data were preprocessed using the fMRIPrep (Esteban et al. 2019) and xcpEngine (Ciric et al. 2017) toolboxes (for detailed steps, see SI Materials and Methods). The preprocessed BOLD time series for each subject was projected onto the cortical surface and smoothed along the surface using a 6mm smoothing kernel using the Freesurfer software package. For each subject and session, the general task-specific activity (convolved with a hemodynamic response function) was regressed out from the BOLD time series, and the time series from all tasks were concatenated into one timeseries per subject and condition.

### Network and assembly identification

For all analyses of task data, the BOLD time series for each vertex on the fsaverage surface was assigned to one of 400 cortical parcels from a pre-determined standardised template of functionally similar cortical regions (Schaefer et al. 2018). The time series for all the vertices corresponding to each parcel were averaged to give one mean time series per parcel, resulting in a total of 400 time series across both cortical hemispheres. We then divided the cortex in three separate steps, which allowed us to arranged the data in a hierarchical framework comprising 4-levels (Fig. S2). Firstly, at the cortex level, each parcel was assigned to either one of 7 functional networks, taken from a template based on a large sample of resting state data (Yeo et al. 2011). At the 2^nd^ level of the hierarchy (7 networks), each parcel was assigned to one of 17 functional networks taken from a higher resolution version of the same template (Yeo et al. 2011), and these 17 networks were clustered into one of the 7 original networks based on naming convention. At the 3^rd^ level (17 networks), each of the 17 functional networks were further partitioned into assemblies (clusters) based upon a hierarchical clustering technique, resulting in a partitioned composed of 57 clusters. To achieve this last level of the hierarchy, the averaged correlation matrix was computed for each network during the RW session and thresholded at p < 0.05. The thresholded correlation structure was computed for each of the 17 networks and their structure was assessed by a hierarchical clustering that maximized intraclass similarity, defined as 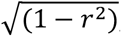, where r is the correlation coefficient between two regions. By thresholding the similarity trees at the level of the highest increase of intracluster distance, the 17 networks were divided into assemblies of areas (Fig. S5). Subcortical regions were extracted from the Freesurfer segmentation of the FSL MNI152 template represented by 8 ROIs corresponding to the left and right hemispherical thalamus, striatum, hippocampus, and amygdala. Partition into community structure and analyses of resting state data are described in SI Materials and Methods.

### Functional connectivity and integration

The mean time course of each parcel or volume was correlated to the mean time courses of all other parcels and the correlation matrices were z-scored. Significance was corrected at p<0.05 using the false discovery rate method (Benjamini et al. 1995).

Functional integration was calculated as follows. The functional data was considered as *N* regions of interest (parcels) characterized by their mean time courses *y* = (*y*_1_ …, *y*_*N*_) gathered into *K* systems (networks) defined as *S* = {*S*_1_ …, *S*_*K*_}. For any partition of *y*, integration (or mutual information) can then defined as the Kullback–Leibler information divergence between the joint distribution *p*(*y*_1_ … *y*_*K*_) and the product of the marginal distributions of the N-dimensional fMRI BOLD time series *y* divided into *K* subsets, such that:

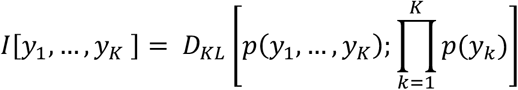

which can be rewritten as:

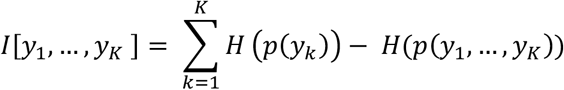

where *H* is the Shannon entropy measure (Marrelec et al. 2008). For multivariate normal data with mean *mu* and covariance matrix *Sigma*, entropy can be computed as:

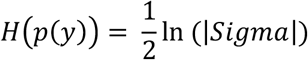

where |*Sigma*| refers to the determinant of the covariance matrix. As described in Marrelec et al (2008), the total integration (***I***_*Tot*_) can be decomposed, according the organization of regions into systems, as the sum of each system’s integration relative to its regions (within-system integration) and a between-system integration term:

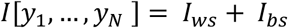

or alternatively:

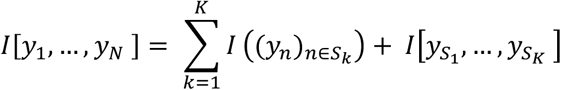

Integration was calculated at different levels in our proposed hierarchical model of the cortex. Firstly, integration was calculated across the whole cortex where N= 400 parcels were gathered into 7 networks. Secondly, for each of 7 networks separately, the N parcels were gathered into small sub-networks (Yeo 17 networks), where each of the 7 networks contained a minimum of 2 out of 17 sub-networks. Finally, for each of the 17 sub-networks, the N parcels were divided into K of the 57 clusters (assemblies), where each of the 17 networks contained a minimum of 2 out of 57 clusters.

We defined the functional clustering ration (FCR) as the ratio of the integration within subsystems (***I***_*ws*,_), compared to the integration present between these subsystems (***I***_*bs*_) (Boly et al. 2012). It is a measure of clustering inside a given system because an increase in FCR indicates that subsystems become proportionally more independent of each other. In other words, it quantifies the degree of functional segregation of a given system into subsystems:

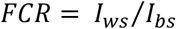

The FCR was also computed for each network or cluster at each level of the hierarchy.

### EEG Data Acquisition and Analysis

EEG was acquired using an MR compatible 256 high-density geodesic sensor EEG array (Electrical Geodesics Inc, *Magstim EGI)*. EEG data were recorded at 1000Hz referenced to Cz using a battery-powered MR-compatible 256-channel amplifier. Electrocardiography (ECG) was also collected via two MR compatible electrodes through a bipolar amplifier (Physiobox, EGI). The EEG data were corrected for MR gradient ballistocardiographic pulse-related artefacts using the Brainvision Analyzer (*Brain Products Inc, Gilching Germany*). The MR-denoised EEG signal was bandpass filtered between 1 and 20 Hz to remove low-frequency drift and high-frequency noise, down-sampled to 250 Hz, and re-referenced to the linked mastoids. Scoring of the sleep session was performed in conjunction by two trained scorers (NEC, AAP) using the Wonambi toolbox (https://github.com/wonambi-python/wonambi) (using the channels Fz, F3, F4, C3, C4, O1 and O2) in order to obtain measures of sleep including total sleep time, and the duration of sleep stages. For the analysis of wake EEG, eye blink artefacts were removed with ICA using the MNE Python package. The ICA was fit on each EEG timeseries concatenated across tasks for each subject and condition. The number of components was set to 15, with a random seed to ensure the uniformity of components. The resulting ICA components were then compared to EOG electrodes any ICs that matched the EOG pattern were automatically marked for exclusion. Any matching components were then excluded from the original signal, and this new timeseries was then used for all subsequent analysis. Power spectral analysis was also computed using MNE, via the Welch method using a frequency resolution of 1Hz.

### Statistical Analysis

#### Bayesian Sampling Scheme

Probable values of integration and FCR were inferred from the data using a Bayesian numerical sampling scheme that approximates the posterior distribution of the parameters of interest (Marrelec et al. 2008). We first ran Gibbs sampling on the model to propose a numerical approximation of *p*(*Sigma*|*y*). Using this sampling procedure, we obtained 1,000 samples from the posterior distribution of the group covariance matrix (*Sigma*_s=1,…,1000_) in either condition (RW, SD, PRN). From each sample *Sigma*_s_, we computed the corresponding values of integration (I_ws_ and I_bs_) and FCR. For each measure and condition, we therefore obtained a frequency histogram that, by construction, approximated the posterior distribution of that measure given the data. The samples were then used to provide approximations of either statistics (e.g., mean and SD) or probabilities (e.g., probability of an increase between RW and SD) approximated as the frequency of that increase observed in the samples. This also allowed to approximate the posterior probability p(A|y) of the assertion e.g. FCR_SD_ > FCR_RW_ using the equation:

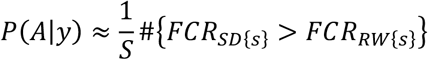

where # stands for the cardinal function of a set. For analyses between conditions, a probability of difference >0.95 was considered significant.

During the sampling procedure, a covariance matrix was also estimated for each individual. Thus, similarly to the group-level analysis, we compared the resulting estimates of FCR at the individual level with other individual metrics (e.g. performance measures, global signal fluctuation, thalamocortical connectivity) using product-moment correlations.

#### Spatial Correspondence Between Cortical Maps

For the comparison between brain maps (Fig 3) we used a spatial permutation framework to generate null models of overlap (Alexander-Bloch et al. 2018). Spatial permutation of brain maps was performed using 100 angular permutations of spherical projections of the cortical surface. This approach calculates a correspondence statistic between two maps while controlling for spatial autocorrelation in the data.

All of the code used in this study is available online at: https://github.com/nathanecross/sleep-deprivation/tree/master/Integration

## Supporting information

Supplementary Methods and Results

## Acknowledgments

This research was supported by the Natural Sciences and Engineering Research Council of Canada (TDV) and the Canada Foundation for Innovation (TDV). The MRI compatible high-density EEG device (Philips Neuro) and data acquisition were made possible through an internal grant from PERFORM center and the Faculty of Arts and Science of Concordia University (CG). TDV is also supported by the Canadian Institutes of Health Research (MOP 142191, PJT 153115, PJT 156125 and PJT 166167), the Fonds de Recherche du Québec – Santé and Concordia University. CG is supported Natural Sciences and Engineering Research Council of Canada Discovery grants as well as the Canadian Institutes of Health Research (PJT-159948 and MOP-133619) and the Fonds de Recherche du Québec, Nature et Technology (research team grant).

