## Supplementary Methods and Results for "An altered balance of integrated and segregated brain activity is a marker of cognitive deficits following sleep deprivation"

**ASI Materials and Methods**

**Population and Experimental Design.**

***Participants***. Participants were recruited using advertisements posted online and within Concordia University, Montreal. A semi-structured interview was conducted to assess their eligibility. Participants were required to be aged between 18 to 30 years and considered good sleepers (>6 hours of sleep per night) with an absence of any sleep disorders (participants with insomnia, sleep apnea syndrome with an apnea-hypopnea index >5/hr, central disorders of hypersomnolence, restless legs syndrome, periodic limb movements during sleep with an index >15/hr, and parasomnias were excluded). Participants were also excluded for neurological or psychiatric conditions (e.g. epilepsy, migraine, stroke, chronic pain, major depression, anxiety disorder, psychotic disorder) and current use of psychotropic medications or cannabis. Subjects were also asked to refrain from smoking tobacco for the entire duration of the study. All subjects provided informed consent prior to the start of the study that was approved by the Central Research Ethics Committee of the Quebec Ministry of Health and Social Services.

***Study Procedure.*** The study protocol is outlined in Fig. S1. Participants made three visits to the laboratory. On the first visit, they were briefed on the study protocol and completed an overnight polysomnography to adapt to the environment and to screen for any sleep disorders. To monitor their sleep pattern between this visit and the remaining visits, each participant was given a wrist actiwatch (Actiwatch, Philips Respironics, USA) that had to be worn until they completed the experiment. Only subjects with a good sleep habit (slept for > 6 hr per night with a consistent schedule) were invited to participate in the subsequent sessions.

The second and third visits were well-rested (WR) or sleep deprivation (SD) sessions. The WR and SD sessions were separated by a minimum of one week and the order of test sessions was counterbalanced across participants. During the WR session, participants arrived at 7:00pm and were given a 9-hour sleep opportunity in a dark, quiet room. During the SD session, participants arrived at 8:00 pm and stayed awake the entire night in the laboratory. They were free to do as they wish, but were constantly monitored and prevented from consuming any stimulants or performing anything too stimulating (e.g. exercising or watching horror films). Light levels were set at a constant level for all subjects throughout the entire night.

In both sessions, subjects were taken to the MRI at 7:00am to be prepared for scanning, including the application of an MRI compatible high-density EEG cap for objective monitoring of wakefulness. Scanning began at 8:00am. In the WR state, the session involved a resting state sequence (a fixation cross (5 minutes), followed by three cognitive tasks: the Attentional Network Task (ANT), the N-back task, and the Mackworth Clock Task (MCT); an anatomical T1w scan, and a diffusion weighted imaging scan. In the SD state, the first session involved the same resting state sequence, followed by the three cognitive tasks. Then subjects were provided a 60-minute nap opportunity inside the MRI scanner. After 60 minutes, the participant was woken and during the final session (post recovery nap, PRN), subjects re-completed all of the cognitive tasks, as well as one resting state sequence (fixation cross). At the completion of the study, subjects were debriefed and were advised not to drive home if they had just completed the SD session.

**Behavioural Data Acquisiton and Analysis.**

The cognitive tasks (Fig. S2) were chosen to reflect different types of cognitive processing due to the known sigificant effects of sleep deprivation in these cognitive domains. All tasks were run on a laptop computer using Inquisit software *(Millisecond Software LLC, Seattle USA*), displayed to the participant via a projector screen behind the MRI scanner. The participant responded to all tasks via button presses made using a response pad attached to the fingers of the left hand. For all tasks, outcomes of reaction time (ms), accuracy (%; correct trials/number of trials), and lapses (number of missed responses) were measured.

***Mackworth Clock Task (MCT)***. The version of this vigilance task used in this study consisted of a circular stimuli presented on screen spatially similar to a ticking clock. A node sequentially traversed the circumference of the circle in discrete steps. The participants were asked to press the trigger when the stimuli skipped or 'jumped' a step (Lichstein et al. 2000, Loh et al. 2004). The task lasted 5 minutes in duration, with each block lasting 18 seconds and rest period between blocks was 15 seconds. Therefore, this task resembles a simple attention task.

***Attentional Network Task (ANT)***. This task probes different attentional processes, such as alerting, orienting and executive control (Fan et al. 2005, Fan et al. 2002). The ANT consists of a series of trials in which the participant is required to identify the direction (left or right) of the middle arrow in an array of five arrows within an upper or lower panel of the screen. The arrow is either congruent or incongruent to the direction of all other arrows. Different cues are displayed immediately prior to the appearance of the arrows, including both panels flashing (double cue; alerting), or either panel flashing (valid or invalid cues; orienting). For a subset of trials, no cue precedes the appearance of the arrows (no cue). The ANT lasted 13 minutes in duration, with each trial lasting 1 seconds with a random jitter (range = 2-12 seconds, mean = 5 seconds) between each trial.

***N-back Task.*** This working memory task consisted of a sequence of letters presented one at a time in the centre of the screen (Kirchner 1958, Sweet 2011). The participant was required to respond depending on the difficulty level of each block of trials. At level 0-back, the participant responded on every trial in the block. This served as a control condition for investigating activations related to higher level processing. At level 1-back, the participant responded whenever the letter of two subsequent trial was the same. At level 2-back, the participant responded whenever the letter presented was the same as the letter presented 2 trials previously. The N-back lasted 8 minutes in duration, with each block lasting 38 seconds with 10 seconds between each trial.

**Monitoring of Wakefulness and Sleep.**

Participants were monitored for wakefulness during scanning through a live video recording of the eyes. They were given a wake-up call if their eyes were closed for more than 10s during the resting-state or cognitive tasks to prevent them from falling asleep. When there was uncertainty surrounding wakefulness inside the scanner, post-hoc confirmation of wakefulness and sleep was verified using the electroencephalography (EEG) that was acquired simultaneously.

**EEG Data Acquisition and Analysis.**

***Acquisition*****.** EEG was acquired using an MR compatible 256 high-density geodesic sensor EEG array (Electrical Geodesics Inc (*EGI),* now *MagStim EGI, Oregon USA*). The EEG cap included 256 sponge electrodes referenced to Cz that covered the entire scalp and part of the face. EEG data were recorded using a battery-powered MR-compatible 256-channel amplifier shielded from the MR environment that was placed next to the participant inside the scanning room. The impedance of the electrodes were initally maintained below 20k$\Omega$ and kept to a maximum of 70k$\Omega$ throughout the recording. Data were sampled at 1000Hz and transferred outside the scanner room through fiber-optic cables to a computer running the Netstation software (*v5, EGI*). The recording of EEG was phase-synchronized to the MR scanner clock (*Sync Clock box*, *EGI*), and all scanner repetition times (TRs) and participant responses were recorded in the EEG traces. Electrocardiography (ECG) was also collected via two MR compatible electrodes placed between the 5$^{th}$ and 7$^{th}$ ribs and above the heart close to the sternum, and recorded at 1000Hz through a bipolar amplifier (*Physiobox, EGI*).

***Preprocessing.*** The EEG data were preprocessed using the Brainvision Analyzer (*Brain Products Inc, Gilching Germany*). Firstly, the EEG data were corrected for MR gradient artefacts using a 21 TRs sliding window template, , which were subtracted from each occurrence of the respective artefacts for each electrode {Allen, 2000 #166}. Ballistocardiographic pulse-related artefacts were separated from the signal using a semi-automatic template time-locked to the detected QRS peaks in the ECG channel and then removed from the EEG signal. The MR-denoised EEG signal was bandpass filtered between 1 and 20 Hz to remove low-frequency drift and high-frequency noise, down-sampled to 250 Hz, and re-referenced to the linked mastoids.

***Scoring.*** The task- and resting-wake EEG recordings were scored post-hoc to confirm participant wakefulness during these sessions. Scoring of the sleep session was performed in conjunction by two trained scorers (NEC, AAP) using the Wonambi toolbox (https://github.com/wonambi-python/wonambi) (using the channels Fz, F3, F4, C3, C4, O1 and O2) in order to obtain measures of sleep including total sleep time (TST), and the duration of sleep stages.

**fMRI Data Acquisition and Analysis.**

***Acquisition.*** MRI scanning was acquired with a 3T GE scanner (General Electric Medical Systems, Wisconsin, US) using an 8-channel head coil. Functional scans were all acquired using a gradient-echo echo-planar imaging (EPI) sequence (TR = 2500ms, TE = 26ms, FA = 90°, 41 transverse slices, 4-mm slice thickness with a 0% inter-slice gap, FOV = 192 x 192 mm, voxel size = 4 x 4 x 4mm$^3$ and matrix size = 64 × 64). High-resolution T1-weighted structural images were acquired using a 3D BRAVO sequence (TR = 7908 ms, TI = 450 ms, TE = 3.06 ms, FA = 12°, 200 slices, voxel size = 1.0 × 1.0 × 1.0 mm, FOV = 256 × 256 mm). During all EEG-fMRI sessions, the helium pump was switched off in order to reduce noise artefacts infiltrating the EEG signal. To minimize movement-related artefacts during the scaning, MRI-compatible foam cushions were used to fix the participant's head in the head coil.

***Preprocessing.*** Preprocessing of fMRI data was performed using fMRIPrep 1.3.1 (Esteban et al. 2019); RRID:SCR_016216), which is based on Nipype 1.1.9 (Esteban et al. 2020, Gorgolewski et al. 2011); RRID:SCR_002502). Firstly, the T1-weighted (T1w) image was corrected for intensity non-uniformity (*ANTs v2.2.0*) and used as T1w-reference throughout the workflow. Then a reference volume of the EPI image and its skull-stripped version were generated using a custom methodology of fMRIPrep. The BOLD reference was then co-registered to the T1w reference using bbregister (*FreeSurfer v6.0*) which implements boundary-based registration (Greve et al. 2009). Co-registration was configured with nine degrees of freedom to account for distortions remaining in the BOLD reference. Head-motion parameters with respect to the BOLD reference (transformation matrices, together with six corresponding rotation and translation parameters) were estimated (*mcflirt*, *FSL v5.0.9; (Jenkinson et al. 2002)* before slice-timing correction (*AFNI, (Cox 1996)*). The BOLD time-series (including slice-timing correction when applied) were resampled onto their original, native space by applying a single, composite transform to correct for head-motion and susceptibility distortions. Transforms were concatenated and applied all at once, with one interpolation (Lanczos) step, so as little information as possible was lost. Frame Displacement and the spatial standard deviation of successive difference images (DVARS) were calculated for each functional run, both using their implementations in Nipype. Brain tissue segmentation of cerebrospinal fluid (CSF), white-matter (WM) and gray-matter (GM) was performed on the brain-extracted T1w using FAST (*FSL v5.0.9*, RRID:SCR_002823, (Zhang et al. 2001). The average timeseries of the cerebrospinal fluid (CSF), white matter (WM), and whole-brain were also extracted. After sampling the BOLD time-series onto their original, native space, these were then resampled via nonlinear transformation to the MNI152NLin2009cAsym standard volumetric space for all subjects for subsequent processing and analysis, keeping the original resolution of the BOLD data.

The BOLD data timeseries in standard space were further denoised using a 36-parameter stream of the xcpEngine (Ciric et al. 2017). First, a temporal filter (0.01-0.08Hz) was applied to the data. Then, six realignment parameters, the mean WM and CSF time series (extracted from fMRIPrep), as well as derivative and quadratic expansions, were all regressed out from the BOLD timeseries.

The final preprocessed BOLD time series for each subject was projected onto the cortical surface (white matter boundary, default) using the fsaverage5 template from the Freesurfer package (*mri_vol2surf; FreeSurfer v6.0*), and smoothed along the surface space using a 6mm smoothing kernel. All MRI preprocessing was performed on the Compute Canada servers Cedar and Graham (https://www.computecanada.ca/research-portal/accessing-resources/available-resources/).

For each subject and session, the preprocessed BOLD timeseries were first concatenated across all tasks, as previously implemented for studying the underlying core of general cognitive processing in the human brain (Elliott et al. 2019, Shine et al. 2019, Zhu et al. 2017). This resulted in a total timeseries of 26 minutes (624 TRs) per session (WR, SD & PRN). The general HRF task-specific activity (i.e. for all trials for the ANT and all blocks for the MCT and N-back tasks) was regressed out from the BOLD timeseries to reduce the influence of task-evoked coactivation in the connectivity analyses (De Havas et al. 2012, Wang et al. 2016). Such an approach has been shown to be comparable to resting state functional connectivity studies and used to extract information about the intrinsic properties of cortical connectivity (Elliott et al. 2019). From each scan sequence aquired during the recovery nap, a continuous period of 26 minutes (i.e. the same length as the concatenated task timeseries, 624 TRs) consisting of stable NREM sleep (i.e. without awakening) was extracted for further analysis for all subjects who managed to sleep inside the scanner (n = 18). These segments were extracted from the earliest periods of continuous (>26min) NREM sleep in each session.

***Network and Assembly Identification.***

For the resting state analyses, these sequences were much shorter (120 TRs) than the concatenated task timeseries (624 TRs). In this case, a 400 parcellation template would not result in a semi-positive definite covariance matrix, as the spatial dimension (400) would be larger than the temporal dimension (120). Therefore, for these analyses we instead used a 100 parcellation template (Schaefer et al. 2018). This reduced number of parcels limited the number of levels in the hierarchy that we could analyse, which is why the resting state analyses are depicted for the cortex and 7 networks only.

To ensure our findings were robust to the use of a pre-defined network template, we replicated the analyses at the whole cortex level using an alternative, data-driven approach taken from graph theory. Specifically, we identified the community structure of cortical functional connectivity using the Louvain modularity algorithm from the Brain Connectivity Toolbox (BCT; {Rubinov, 2011 #170}). Community structure was identified from the signed, weighted 400x400 correlation matrix of the concatenated task-regressed BOLD time series during the WR condition. Using these community assignments, integration and FCR were computed using the same technique as done for the Yeo network templates.

***Functional Connectivity.*** The mean time course of each parcel (400 for task, 100 for resting-state) was correlated with the mean time course of every other parcel for each session (state) and the correlation matrices (in each session) were z-scored to encourage normality. Repeated measures t-tests were performed for each parcel between the RW and SD state. For the change between the SD and PRN state, a repeated-measures ANOVA was computed, including the total sleep time as a covariate. Significance was corrected at $p<0.05$ using the false discovery rate method (Benjamini et al. 1995).

***References***

Alexander-Bloch, A. F., H. Shou, S. Liu, T. D. Satterthwaite, D. C. Glahn, R. T. Shinohara, S. N. Vandekar and A. Raznahan On testing for spatial correspondence between maps of human brain structure and function. NeuroImage. 2018; **178**: 540-551.
Benjamini, Y. and Y. Hochberg Controlling the False Discovery Rate: A Practical and Powerful Approach to Multiple Testing. Journal of the Royal Statistical Society. Series B (Methodological). 1995; **57**(1): 289-300.
Boly, M., V. Perlbarg, G. Marrelec, M. Schabus, S. Laureys, J. Doyon, M. Pélégrini-Issac, P. Maquet and H. Benali Hierarchical clustering of brain activity during human nonrapid eye movement sleep. Proceedings of the National Academy of Sciences of the United States of America. 2012; **109**(15): 5856-5861.
Ciric, R., D. H. Wolf, J. D. Power, D. R. Roalf, G. L. Baum, K. Ruparel, R. T. Shinohara, M. A. Elliott, S. B. Eickhoff, C. Davatzikos, R. C. Gur, R. E. Gur, D. S. Bassett and T. D. Satterthwaite Benchmarking of participant-level confound regression strategies for the control of motion artifact in studies of functional connectivity. Neuroimage. 2017; **154**: 174-187.
Cox, R. W. AFNI: software for analysis and visualization of functional magnetic resonance neuroimages. Comput Biomed Res. 1996; **29**(3): 162-173.
De Havas, J. A., S. Parimal, C. S. Soon and M. W. L. Chee Sleep deprivation reduces default mode network connectivity and anti-correlation during rest and task performance. NeuroImage. 2012; **59**: 1745-1751.
Elliott, M. L., A. R. Knodt, M. Cooke, M. J. Kim, T. R. Melzer, R. Keenan, D. Ireland, S. Ramrakha, R. Poulton, A. Caspi, T. E. Moffitt and A. R. Hariri General functional connectivity: Shared features of resting-state and task fMRI drive reliable and heritable individual differences in functional brain networks. Neuroimage. 2019; **189**: 516-532.
Esteban, O., C. J. Markiewicz, R. W. Blair, C. A. Moodie, A. I. Isik, A. Erramuzpe, J. D. Kent, M. Goncalves, E. DuPre, M. Snyder, H. Oya, S. S. Ghosh, J. Wright, J. Durnez, R. A. Poldrack and K. J. Gorgolewski fMRIPrep: a robust preprocessing pipeline for functional MRI. Nat Methods. 2019; **16**(1): 111-116.
Esteban, O., C. J. Markiewicz, H. Johnson, E. Ziegler, A. Manhes-Savio, D. Jarecka, C. Burns, D. G. Ellis, C. Hamalainen, M. P. Notter, B. Yvernault, T. Salo, M. Waskom, M. Goncalves, K. Jordan, J. Wong, B. E. Dewey, C. Madison, E. Benderoff, D. Clark, F. Loney, D. Clark, A. Keshavan, M. Joseph, D. M. Nielson, M. Dayan, M. Modat, A. Gramfort, S. Bougacha, B. Pinsard, S. Berleant, H. Christian and A. a. Rokem nipy/nipype: 1.4.2. 2020.
Fan, J., B. D. McCandliss, J. Fossella, J. I. Flombaum and M. I. Posner The activation of attentional networks. Neuroimage. 2005; **26**(2): 471-479.
Fan, J., B. D. McCandliss, T. Sommer, A. Raz and M. I. Posner Testing the efficiency and independence of attentional networks. J Cogn Neurosci. 2002; **14**(3): 340-347.
Gorgolewski, K., C. D. Burns, C. Madison, D. Clark, Y. O. Halchenko, M. L. Waskom and S. S. Ghosh Nipype: a flexible, lightweight and extensible neuroimaging data processing framework in python. Front Neuroinform. 2011; **5**: 13.
Greve, D. N. and B. Fischl Accurate and robust brain image alignment using boundary-based registration. Neuroimage. 2009; **48**(1): 63-72.
Jenkinson, M., P. Bannister, M. Brady and S. Smith Improved optimization for the robust and accurate linear registration and motion correction of brain images. Neuroimage. 2002; **17**(2): 825-841.
Kirchner, W. K. Age differences in short-term retention of rapidly changing information. J Exp Psychol. 1958; **55**(4): 352-358.
Lichstein, K. L., B. W. Riedel and S. L. Richman The Mackworth Clock Test: a computerized version. J Psychol. 2000; **134**(2): 153-161.
Loh, S., N. Lamond, J. Dorrian, G. Roach and D. Dawson The validity of psychomotor vigilance tasks of less than 10-minute duration. Behav Res Methods Instrum Comput. 2004; **36**(2): 339-346.
Marrelec, G., P. Bellec, A. Krainik, H. Duffau, M. Pélégrini-Issac, S. Lehéricy, H. Benali and J. Doyon Regions, systems, and the brain: hierarchical measures of functional integration in fMRI. Med Image Anal. 2008; **12**(4): 484-496.

Millisecond Software LLC. (, Seattle USA).
Schaefer, A., R. Kong, E. M. Gordon, T. O. Laumann, X. N. Zuo, A. J. Holmes, S. B. Eickhoff and B. T. T. Yeo Local-Global Parcellation of the Human Cerebral Cortex from Intrinsic Functional Connectivity MRI. Cereb Cortex. 2018; **28**(9): 3095-3114.
Shine, J. M., M. Breakspear, P. T. Bell, K. A. Ehgoetz Martens, R. Shine, O. Koyejo, O. Sporns and R. A. Poldrack Human cognition involves the dynamic integration of neural activity and neuromodulatory systems. Nat Neurosci. 2019; **22**(2): 289-296.
Sweet, L. H. N-Back Paradigm. Encyclopedia of Clinical Neuropsychology. 2011 J. S. Kreutzer, J. DeLuca and B. Caplan. New York, NY, Springer New York**:** 1718-1719.
Wang, C., J. L. Ong, A. Patanaik, J. Zhou and M. W. Chee Spontaneous eyelid closures link vigilance fluctuation with fMRI dynamic connectivity states. Proc Natl Acad Sci U S A. 2016; **113**(34): 9653-9658.
Yeo, B. T., F. M. Krienen, J. Sepulcre, M. R. Sabuncu, D. Lashkari, M. Hollinshead, J. L. Roffman, J. W. Smoller, L. Zollei, J. R. Polimeni, B. Fischl, H. Liu and R. L. Buckner The organization of the human cerebral cortex estimated by intrinsic functional connectivity. J Neurophysiol. 2011; **106**(3): 1125-1165.
Zhang, Y., M. Brady and S. Smith Segmentation of brain MR images through a hidden Markov random field model and the expectation-maximization algorithm. IEEE Trans Med Imaging. 2001; **20**(1): 45-57.
Zhu, Y., L. Cheng, N. He, Y. Yang, H. Ling, H. Ayaz, S. Tong, J. Sun and Y. Fu Comparison of Functional Connectivity Estimated from Concatenated Task-State Data from Block-Design Paradigm with That of Continuous Task. Computational and mathematical methods in medicine. 2017; **2017**: 4198430-4198430.

**SI Results**

**Behavioural Results.**

Mean performance scores during each task are depicted in Fig. S1 and Table S1. As expected, across all tasks outcomes were significantly impaired following sleep deprivation, and improved following the recovery nap. In the MCT, mean reaction time increased from the WR to the SD state (q=5.41, 95%CI = -43.6 to -8.8, p=0.003), and decreased from the SD to PRN state (q= -5.82, 95%CI = 11.2 to 47.3, p=0.004), but there were no differences between the WR and PRN states (q=0.62, 95%CI = -14.5 to 20.5, p=0.673; $F_{1,19}$ = 10.71, $\eta^2$ = 0.056; p<0.001). Accuracy on the MCT also decreased from the WR to the SD state (q=-4.73, 95%CI = 2.8 to 20.9, p=0.009), and increased from the SD to PRN state (q = 4.59, 95%CI = -19.2 to -2.3, p=0.011), but there were no differences between the WR and PRN states (q=0.63, 95%CI = -5.5 to 7.8, p=0.606; $F_{1,19}$ =8.45, $\eta^2$ = 0.078, p=0.002). In the ANT, there were mean reaction time differences between the WR and SD states (q=5.66, 95%CI = -111.4 to -24.9, p=0.002), and between the SD and PRN states (q= -5.45, 95%CI = 18.9 to 91.9, p=0.003), but not between the WR and PRN states (q=1.26, 95%CI = -49.3 to 23.8, p=0.654; $F_{1,19}$ =11.19, $\eta^2$ = 0.093, p<0.001). Accuracy decreased from the WR to the SD state (q=5.43, 95%CI = 7.1 to 34.6, p=0.003), and decreased from the SD to PRN state (q= -5.03, 95%CI = -25.4 to -4.2, p=0.006), but there were no differences between the WR and PRN states (q=2.13, 95%CI = -4.1 to 16.2, p=0.309; $F_{1,19}$ = 10.99, $\eta^2$ =0.204, p<0.001). In the 2-back task, mean reaction time increased between the WR and SD states (q=5.31, 95%CI = -366.1 to -70.6, p=0.004), and decreased between the SD and PRN states (q= -4.42, 95%CI = 27.5 to 267.0, p=0.015), but did not significantly change between the WR and PRN states (q=2.78, 95%CI = -163.1 to 20.8, p=0.148; $F_{1,19}$ = 10.76, $\eta^2$ = 0.161, p<0.001). Accuracy decreased from the WR to the SD state (q=5.02, 95%CI = 3.2 to 19.5, p=0.005), and decreased from the SD to PRN state (q= -4.73, 95%CI = -13.7 to -1.9, p=0.009), but there were no differences between the WR and PRN states (q=2.88, 95%CI = -0.9 to 8.1, p=0.13; $F_{1,19}$ = 10.77, $\eta^2$ =0.226, p=0.001).

**Changes in Resting-State Functional Integration**

In the resting state sequence, there was an increase in FCR between RW and SD. $I_{Tot}$ also increased between the RW and SD resting state. Integration increased within all of the 7 networks (Fig. S3). Following the PRN the resting-state FCR significantly decreased across the cortex and within all of the 7 networks.

**Integration changes during sleep**

There was significantly greater variance in the magnitude of covariance between time series of almost all edges time series in the NREM condition compared to the WR state (90% of edges passing significance for Levene's test p<0.05). As such, group-inference for this condition was not applicable because this increased variability of covariance led to higher estimates of between- and within-integration at the group level than any individual exhibited alone. Thus, for comparisons between the NREM condition and other conditions, significance of integration was conducted via Friedman's test and post-hoc Wilcoxon rank sum tests between estimates of FCR and integration taken from individual covariance matrices. There was a significant effect of condition on FCR ($\chi^2$ = 41.35, p=0.0008). Post-hoc tests revealed significant differences between WR and NREM after correction for multiple comparisons (Z = -5.11; $p_{FDR}$ <0.001), SD and NREM (Z = -4.64; $p_{FDR}$ <0.001 ), and PRN and NREM (Z = -4.86; $p_{FDR}$ <0.001). In summary, the FCR was highest in the NREM condition compared to other conditions (Fig 2B), and this was driven by both an increase in within-systems integration as well as a decrease in between-systems integration (Fig. S4). $I_{Tot}$ was also significantly different across conditions ($\chi^2$ = 42.25, p=0.0006). However, post-hoc tests revealed that none of differences between NREM and wake conditions passed the threshold for significance after correction for multiple comparisons (all $p_{FDR}$ >0.05).

The change in FCR from SD to the PRN was not related to the duration of total sleep time (TST; r = 0.12, p = 0.620) or percentage of NREM Stage 3 (r = -0.30, p = 0.207). Neither was the change in $I_{Tot}$

from SD to the PRN related to the duration of total sleep time (TST; r = 0.09, p = 0.712) or percentage of NREM Stage 3 (r = -0.14, p = 0.547).

**Changes in Functional Integration Using Networks Defined By Community Structure**

5 networks (communities) were detected from the functional connectome during the WR condition using the Louvain modularity algorithm (Fig. S9a). Integration within and between these networks also increased between RW and SD (Fig. S9b), and decreased (but of a lesser magnitude) from the SD to the PRN session (Table S6).

**Relationships Between Integration and Performance.**

The first component for Accuracy performance was robustly and significantly negatively associated with the change in $I_{Tot}$ (r = -0.87, p<0.001; Fig. S6). The first component for Speed was also significantly positively correlated with the change in $I_{Tot}$ (r = 0.69, p < 0.001). Linear rank (Spearman's) correlations revealed that these correlations remained significant: Accuracy and $I_{Tot}$ (rho = -0.71, p <0.0001; Reaction Time and $I_{Tot}$ (rho = 0.52, p = 0.019).We also assessed the change in performance on each task separately in comparison to the change in integration and FCR of cortical BOLD activity during that task only (Fig. S7). The extent of the increase in FCR within each individual was negatively correlated with change in performance accuracy from baseline to SD on the ANT (r = -0.66, $p_{FDR}$ = 0.004) and N-back (r = -0.61, $p_{FDR}$ = 0.011) tasks, but not the MCT (r = -0.01, $p_{FDR}$ = 0.961). The change in FCR was also positively correlated with change in mean reaction time during the ANT (r = 0.53, $p_{FDR}$ = 0.027) and N-back task (r=0.58, $p_{FDR}$ = 0.015), but not the MCT (r = 0.14, $p_{FDR}$ = 0.663). The extent of the increase in $I_{Tot}$ within each individual was negatively correlated with change in performance accuracy from baseline to SD on both the ANT (r = -0.83, $p_{FDR}$<0.001) and N-back (r = -0.70, $p_{FDR}$ = 0.003) tasks, but not the MCT (r = -0.07, $p_{FDR}$ = 0.852). The change in $I_{Tot}$was also positively correlated with change in mean reaction time during the ANT (r = 0.52, $p_{FDR}$ = 0.027) and the N-back task (r = 0.73, $p_{FDR}$ = 0.002), but not the MCT (r = 0.17, $p_{FDR}$ = 0.634).

On the network level, the change in integration within each of the 7 networks was negatively correlated with change in Accuracy from WR to SD: Visual (r = -0.78, $p_{FDR}$ < 0.001), Somatomotor (r = -0.72, $p_{FDR}$ < 0.001), Dorsal Attention (r = -0.84, $p_{FDR}$ < 0.001), Ventral Attention (r = -0.73, $p_{FDR}$ < 0.001), Limbic (r = -0.50, $p_{FDR}$ = 0.026), Default Mode (r = -0.81, $p_{FDR}$ < 0.001), and Frontoparietal network (r = -0.79, $p_{FDR}$ < 0.001). Results were similar for Speed performance: Visual (r = -0.78, $p_{FDR}$ < 0.001), Somatomotor (r = -0.72, $p_{FDR}$ < 0.001), Dorsal Attention (r = -0.84, $p_{FDR}$ < 0.001), Ventral Attention (r = -0.73, $p_{FDR}$ < 0.001), Limbic (r = -0.50, $p_{FDR}$ = 0.026), Default Mode (r = -0.81, $p_{FDR}$ < 0.001), and Frontoparietal network (r = -0.79, $p_{FDR}$ < 0.001).

**Relationships Between Integration and Global Signal Fluctuations**

The global signal fluctuation was significantly associated with average frame displacement (FD) in each session (WR: r = 0.76, p < 0.001; SD: r = 0.58, p = 0.008; PRN: r = 0.66, p = 0.001). However, the FCR and FD were not correlated in any vigilance state (WR: r = 0.13, p = 0.598; SD: r = 0.31, p = 0.189; PRN: r = -0.06, p = 0.801). Furthermore, the relationship between the global signal fluctuation and FCR was still significantly correlated, even after including average FD in each state as a covariate in statistical models (WR: r = 0.76, p < 0.001; SD: r = 0.79, p < 0.001; PRN: r = 0.67, p = 0.002).

**Relationships Between Integration and Homeostatic Markers of Sleep Pressure**

There were no significant associations between the change in integration or FCR with any marker of sleep homeostatic pressure. Firstly, subjective sleepiness measured via the Karolinska Sleepiness Score (KSS) was not associated with the change from RW to SD in integration (r = -0.04, p = 0.864) or FCR (r = -0.14, p = 0.546). Secondly, the change from RW to SD in the power of EEG frequencies 1-10Hz were not related to the change in total integration (r values = 0.10-0.39, all p>0.129) or FCR (r values = -0.01-0.21, all p>0.413). Neither did the magnitude of the increase in FCR following SD correlate with sleep latency (r = -0.06, p = 0.804), latency to Stage 3 NREM sleep (r = -0.30, p = 0.255) or the percentage of NREM Stage 3 during the nap (r = 0.30, p = 0.198).

**Relationships Between Performance and Homeostatic Markers of Sleep Pressure**

There were no significant associations between the change in general performance on cognitive tasks with any marker of sleep homeostatic pressure. Sleep latency was not associated with change in accuracy score from WR to SD (r = 0.20, p = 0.417) or change in reaction time (r = -0.29, p = 0.224). Latency to NREM3 was not associated with change in accuracy score from WR to SD (r = 0.25, p = 0.346) or change in reaction time (r = -0.09, p = 0.718). Additionally, percentage of NREM Stage 3 during the nap was not associated with change in accuracy score from WR to SD (r = -0.24, p = 0.322) or change in reaction time (r = 0.24, p = 0.327). Finally, the change from RW to SD in the power of EEG frequencies 1-10Hz were not related to the change in accuracy from WR to SD (r values = -0.12-0.13, all p>0.613) or change in reaction time (r values = -0.21-0.13, all p>0.424).

**Changes in Thalamocortical Connectivity**

Despite widespread reductions in thalamocortical functional connectivity (correlation coefficients) from WR to SD, there were no associations between the magnitude of thalamocortical changes and accuracy (r = -0.06, p = 0.804) or speed (r = -0.14, p = 0.540) performance on the cognitive tasks. There were also no significant associations between the change in thalamocortical connectivity with any marker of sleep homeostatic pressure: KSS subjective sleepiness (r = 0.13, p = 0.581), sleep latency (r = 011, p = 0.651), latency to Stage 3 NREM sleep (r = -0.275, p = 0.303) or percentage of NREM Stage 3 during the nap (r = -0.12, p = 0.633).

**Table S1**. Mean task performance scores in each state.

| | | RW | SD | PRN |
|---|---|---|---|---|
| ***PVT*** | Reaction Time (ms) | 473.35 ± 59 | 500.20 ± 46 | 468.9 ± 56 |
| | Accuracy (%) | 72.1 ± 19 | 60.2 ± 16 | 71.1 ± 17 |
| ***ANT*** | Reaction Time (ms) | 736.56 ± 113 | 809.30 ± 131 | 751.31 ± 98 |
| | Accuracy (%) | 89.1 ± 13 | 64.8 ± 22 | 82.0 ± 18 |
| ***N-back*** | Reaction Time (ms) | 699.25 ± 235 | 1003.51 ± 416 | 783.49 ± 249 |
| | Accuracy (%) | 95.8 ± 3 | 84.7 ± 13 | 92.7 ± 7 |

**Table S2**. The $I_{Tot}$ within 7 cortical networks, in the WR, SD and PRN states.

| | $WR_{mean}$ | $WR_{std}$ | p>0.95† | $SD_{mean}$ | $SD_{std}$ | p>0.95† | $PRN_{mean}$ | $PRN_{std}$ |
|---|---|---|---|---|---|---|---|---|
| VN | 11.20 | 0.20 | + | 12.74 | 0.22 | - | 12.09 | 0.22 |
| SMN | 15.62 | 0.21 | + | 17.76 | 0.25 | - | 16.82 | 0.23 |
| DAN | 10.18 | 0.20 | + | 11.89 | 0.22 | - | 11.37 | 0.21 |
| VAN | 8.44 | 0.17 | + | 9.52 | 0.19 | = | 9.82 | 0.20 |
| LIM | 3.00 | 0.12 | = | 3.06 | 0.13 | = | 2.98 | 0.12 |
| FPN | 14.01 | 0.20 | + | 14.49 | 0.21 | = | 15.23 | 0.21 |
| DMN | 14.66 | 0.19 | + | 16.20 | 0.21 | - | 15.28 | 0.20 |

†Approximated posterior probability p(A|y) of the assertion that one state is greater than the other (e.g. $FCR_{SD}$ > $FCR_{WR}$). Calculated by as the frequency of the increase observed in the Bayesian sampling scheme (see Statistical Analysis).

**Table S3**. The $I_{Tot}$ within 17 cortical networks, in the WR, SD and PRN states.

| | $WR_{mean}$ | $WR_{std}$ | p>0.95 | $SD_{mean}$ | $SD_{std}$ | p>0.95 | $PRN_{mean}$ | $PRN_{std}$ |
|---|---|---|---|---|---|---|---|---|
| VisCent | 5.61 | 0.17 | + | 6.74 | 0.20 | = | 6.35 | 0.20 |
| VisPeri | 6.56 | 0.19 | + | 7.44 | 0.23 | - | 6.82 | 0.21 |
| SomMotA | 10.07 | 0.22 | + | 11.82 | 0.24 | = | 11.25 | 0.26 |
| SomMotB | 6.93 | 0.18 | + | 8.31 | 0.20 | - | 7.58 | 0.19 |
| DorsAttnA | 4.28 | 0.14 | + | 4.98 | 0.16 | = | 5.22 | 0.17 |
| DorsAttnB | 4.04 | 0.14 | + | 5.29 | 0.17 | = | 5.12 | 0.16 |
| SalVentAttnA | 5.48 | 0.15 | + | 6.29 | 0.16 | = | 6.08 | 0.16 |
| SalVentAttnB | 2.94 | 0.13 | = | 2.98 | 0.13 | = | 2.99 | 0.13 |
| LimbicA | 1.63 | 0.12 | = | 1.66 | 0.12 | = | 2.03 | 0.15 |
| LimbicB | 2.66 | 0.14 | = | 2.91 | 0.13 | = | 2.84 | 0.14 |
| ContA | 4.05 | 0.14 | = | 4.30 | 0.15 | = | 4.55 | 0.16 |
| ContB | 3.51 | 0.13 | + | 3.96 | 0.14 | = | 4.01 | 0.15 |
| ContC | 2.25 | 0.11 | = | 2.23 | 0.10 | = | 2.25 | 0.11 |
| DefaultA | 7.48 | 0.17 | + | 7.86 | 0.18 | = | 7.49 | 0.18 |
| DefaultB | 5.18 | 0.15 | + | 5.86 | 0.16 | - | 5.46 | 0.16 |
| DefaultC | 2.05 | 0.09 | + | 2.31 | 0.09 | = | 2.17 | 0.10 |
| TempPar | 2.06 | 0.11 | + | 2.91 | 0.14 | = | 2.64 | 0.13 |

†Approximated posterior probability p(A|y) of the assertion that one state is greater than the other (e.g. $FCR_{SD}$ > $FCR_{WR}$). Calculated by as the frequency of the increase observed in the Bayesian sampling scheme (see Statistical Analysis).

**Table S4**. The FCR within 7 cortical networks, in the WR, SD and PRN states.

| | $WR_{mean}$ | $WR_{std}$ | p>0.95 | $SD_{mean}$ | $SD_{std}$ | p>0.95 | $PRN_{mean}$ | $PRN_{std}$ |
|---|---|---|---|---|---|---|---|---|
| VN | 4.16 | 0.10 | = | 4.36 | 0.09 | = | 4.34 | 0.10 |
| SMN | 9.06 | 0.22 | + | 9.67 | 0.21 | = | 9.49 | 0.23 |
| DAN | 8.08 | 0.25 | = | 8.49 | 0.24 | = | 8.20 | 0.24 |
| VAN | 4.79 | 0.15 | + | 5.17 | 0.14 | = | 5.17 | 0.14 |
| LIM | 15.39 | 1.71 | = | 14.32 | 1.71 | = | 15.47 | 1.79 |
| FPN | 2.80 | 0.05 | + | 3.02 | 0.05 | = | 3.03 | 0.06 |
| DMN | 3.56 | 0.07 | + | 3.87 | 0.07 | - | 3.60 | 0.07 |

†Approximated posterior probability p(A|y) of the assertion that one state is greater than the other (e.g. $FCR_{SD}$ > $FCR_{WR}$).

Calculated by as the frequency of the increase observed in the Bayesian sampling scheme (see Statistical Analysis).

**Table S5**. The FCR within 17 cortical networks, in the WR, SD and PRN states.

| | $WR_{mean}$ | $WR_{std}$ | p>0.95 | $SD_{mean}$ | $SD_{std}$ | p>0.95 | $PRN_{mean}$ | $PRN_{std}$ |
|---|---|---|---|---|---|---|---|---|
| ConA | 4.17 | 0.08 | + | 4.01 | 0.08 | = | 4.09 | 0.08 |
| ConB | 6.39 | 0.21 | = | 7.92 | 0.19 | = | 6.91 | 0.20 |
| ConC | 2.81 | 0.12 | = | 3.19 | 0.11 | = | 3.31 | 0.11 |
| TemP | 3.47 | 0.22 | = | 3.41 | 0.20 | = | 3.55 | 0.22 |
| DmnA | 2.83 | 0.22 | = | 2.90 | 0.21 | = | 2.85 | 0.23 |
| DmnB | 4.18 | 0.29 | = | 4.72 | 0.26 | = | 4.17 | 0.28 |
| DmnC | 7.06 | 0.10 | + | 5.30 | 0.10 | = | 6.59 | 0.10 |
| DorsA | 4.84 | 0.16 | = | 5.19 | 0.18 | = | 5.45 | 0.17 |
| DorsB | 5.77 | 0.17 | = | 5.98 | 0.18 | = | 6.07 | 0.17 |
| LimA | 1.70 | 0.03 | = | 1.70 | 0.03 | + | 1.84 | 0.33 |
| LimB | 0.37 | 0.20 | = | 0.39 | 0.18 | = | 1.78 | 0.19 |
| SalA | 2.90 | 0.44 | + | 3.19 | 0.57 | = | 3.10 | 0.47 |
| SalB | 2.82 | 0.18 | = | 3.08 | 0.23 | = | 2.96 | 0.25 |
| SomA | 4.47 | 0.09 | = | 4.64 | 0.09 | = | 4.41 | 0.09 |
| SomB | 5.58 | 0.19 | + | 5.81 | 0.18 | - | 5.48 | 0.18 |
| VisC | 2.23 | 0.68 | = | 2.48 | 0.41 | = | 2.42 | 0.64 |
| VisP | 5.38 | 0.28 | = | 4.95 | 0.21 | = | 5.15 | 0.25 |

†Approximated posterior probability p(A|y) of the assertion that one state is greater than the other (e.g. $FCR_{SD}$ > $FCR_{WR}$). Calculated by as the frequency of the increase observed in the Bayesian sampling scheme (see Statistical Analysis).

**Table S6**. Changes in total integration and FCR across the whole cortex and cortical networks, when task-specific activity was not regressed out of the time series for each parcel.

| Level | $WR_{mean}$ | $WR_{std}$ | p>0.95 | $SD_{mean}$ | $SD_{std}$ | p>0.95 | $PRN_{mean}$ | $PRN_{std}$ |
|---|---|---|---|---|---|---|---|---|
| Cortex | 97.71 | 0.40 | + | 102.06 | 0.41 | - | 100.42 | 0.42 |
| *7 Networks* | | | | | | | | |
| VN | 10.91 | 0.19 | + | 12.36 | 0.21 | - | 11.78 | 0.20 |
| SMN | 16.32 | 0.20 | + | 18.68 | 0.24 | - | 17.95 | 0.24 |
| DAN | 12.27 | 0.19 | + | 13.98 | 0.20 | - | 13.28 | 0.21 |
| VAN | 11.82 | 0.19 | + | 13.49 | 0.21 | - | 12.77 | 0.21 |
| LIM | 5.53 | 0.15 | + | 6.31 | 0.16 | = | 6.01 | 0.17 |
| FPN | 17.58 | 0.21 | + | 19.96 | 0.25 | - | 19.37 | 0.25 |
| DMN | 18.01 | 0.21 | + | 20.33 | 0.23 | = | 19.82 | 0.24 |
| *17 Networks* | | | | | | | | |
| ConA | 6.25 | 0.17 | + | 7.05 | 0.19 | = | 6.71 | 0.19 |
| ConB | 5.89 | 0.17 | + | 6.73 | 0.18 | = | 6.39 | 0.19 |
| ConC | 9.44 | 0.19 | + | 11.19 | 0.20 | - | 10.59 | 0.20 |
| TemP | 7.56 | 0.17 | + | 8.80 | 0.18 | - | 8.33 | 0.19 |
| DmnA | 6.50 | 0.16 | + | 7.37 | 0.18 | = | 7.02 | 0.19 |
| DmnB | 6.27 | 0.17 | + | 7.08 | 0.18 | = | 6.74 | 0.17 |
| DmnC | 8.31 | 0.17 | + | 9.73 | 0.19 | - | 9.25 | 0.18 |
| DorsA | 4.04 | 0.14 | + | 4.71 | 0.15 | = | 4.41 | 0.15 |
| DorsB | 3.14 | 0.14 | + | 3.49 | 0.14 | = | 3.37 | 0.15 |
| LimA | 2.66 | 0.14 | = | 2.91 | 0.13 | = | 2.84 | 0.14 |
| LimB | 6.25 | 0.17 | + | 7.04 | 0.19 | = | 6.72 | 0.18 |
| SalA | 6.28 | 0.17 | + | 7.10 | 0.19 | = | 6.75 | 0.19 |
| SalB | 2.95 | 0.14 | = | 3.28 | 0.15 | = | 3.24 | 0.16 |
| SomA | 8.32 | 0.16 | + | 9.74 | 0.19 | - | 9.25 | 0.19 |
| SomB | 7.72 | 0.17 | + | 8.97 | 0.18 | - | 8.50 | 0.19 |
| VisC | 3.14 | 0.14 | + | 3.50 | 0.14 | = | 3.38 | 0.15 |
| VisP | 3.64 | 0.14 | + | 4.17 | 0.15 | = | 3.96 | 0.15 |

†Approximated posterior probability p(A|y) of the assertion that one state is greater than the other (e.g. $FCR_{SD}$ > $FCR_{WR}$). Calculated by as the frequency of the increase observed in the Bayesian sampling scheme (see Statistical Analysis).

**Table S7**. Changes in total integration and FCR across the whole cortex using networks defined by the Louvain modularity algorithm.

| | $WR_{mean}$ | $WR_{std}$ | p > 0.95 | $SD_{mean}$ | $SD_{std}$ | p > 0.95 | $PRN_{mean}$ | $PRN_{std}$ |
|---|---|---|---|---|---|---|---|---|
| Total Integration | 97.70 | 0.40 | + | 102.10 | 0.40 | - | 100.40 | 0.40 |
| FCR | 1.23 | 0.01 | + | 1.310 | 0.01 | - | 1.27 | 0.01 |

†Approximated posterior probability p(A|y) of the assertion that one state is greater than the other (e.g. $FCR_{SD}$ > $FCR_{WR}$).

Calculated by as the frequency of the increase observed in the Bayesian sampling scheme (see Statistical Analysis).

**Table S8**. Mean sleep statistics during the nap.

| | **Mean** | **Std** |
|---|---|---|
| Sleep Latency, min | 5.32 | 9.77 |
| Total Sleep Time, min | 50.45 | 12.04 |
| Sleep Efficiency, % | 81.83 | 18.98 |
| NREM1, % | 6.67 | 5.11 |
| NREM2, % (n=19) | 72.65 | 18.64 |
| NREM3, % (n=16) | 20.68 | 18.01 |
| Sleep Fragmentation, $n \cdot hr^{-1}$ | 8.91 | 5.62 |
| Latency to NREM3, min | 15.91 | 5.16 |

A| Task performances across states

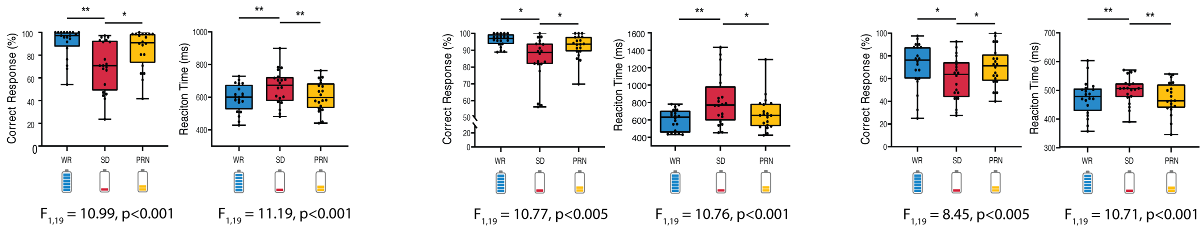


Fig S1. Performance outcomes (accuracy + reaction time) across all tasks were significantly impaired following sleep deprivation, and improved following the recovery nap.

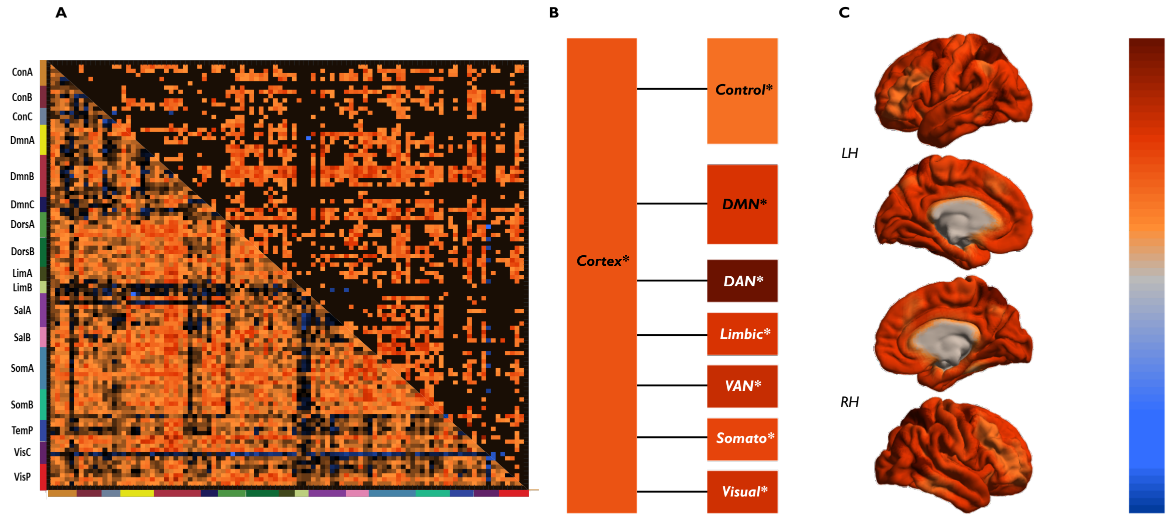


Fig S2 - **(A)** The matrix of change in resting-state functional correlations between the time series of 100 cortical parcels depicts a widespread increase in functional connectivity from well rested (WR) to sleep deprived (SD) states, however to a lesser extent than during tasks (unthresholded, lower triangle; thresholded $P_{FDR}<0.05$, upper triangle). **(B)** Changes in integration are shown across two levels of a hierarchical model of the cortex: whole cortex and 7 networks (the other levels could not be calculated with a 100 parcellation template, due to spatial resolution and that integration calculations required >1 parcels per region). The total integration within the cortex increased from the WR to the SD state, and within 6 out of the 7 functional networks (white italics depict significant changes, variations probability > 0.95). **(C)** The change in total integration mapped onto the cortical surface illustrates a widespread increase in integration across the entire cortex during resting-state.

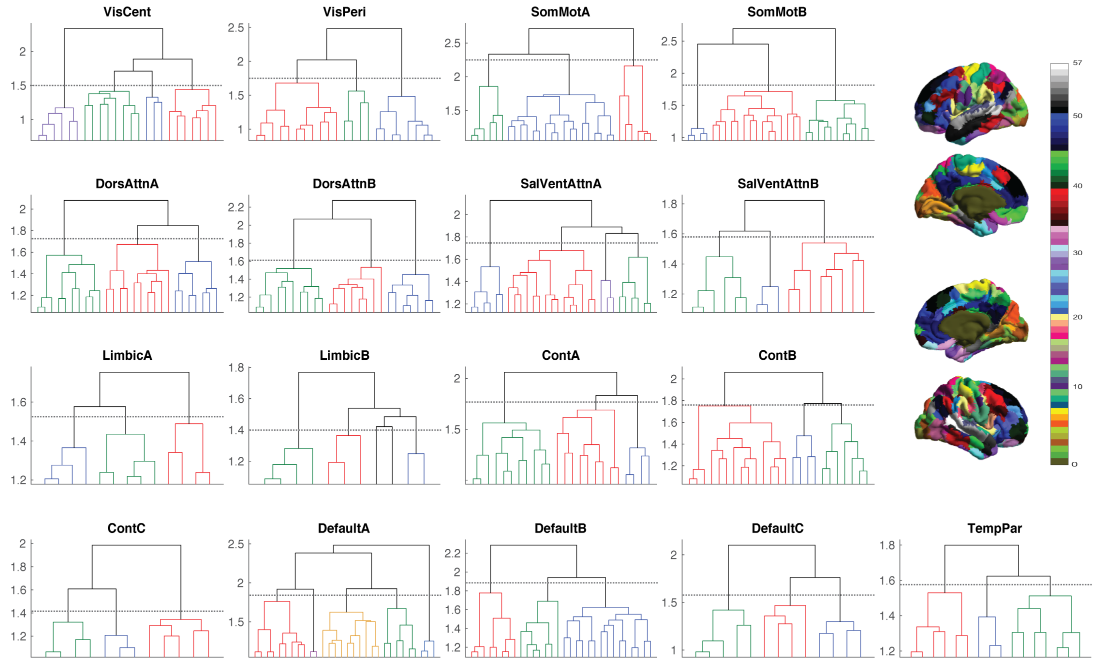


Fig. S3 - Each of the 17 Yeo functional networks were partitioned into assemblies (clusters) based upon a hierarchical clustering that maximized intraclass similarity. A total whole brain partition of 57 clusters was resulting from this procedure. To achieve this, the averaged correlation matrix was computed for each network during the WR session and thresholded at $p < 0.05$. The thresholded correlation structure was computed for each of the 17 networks and their structure was assessed by a hierarchical clustering that maximized intraclass similarity, defined as $\sqrt{(1-r^2)}$, where r is the correlation coefficient between two regions. By thresholding the similarity trees at the level of the highest increase of intracluster distance (dotted line), the 17 networks were divided into assemblies of areas. Clusters are displayed upon the cortical surface (*upper right*).

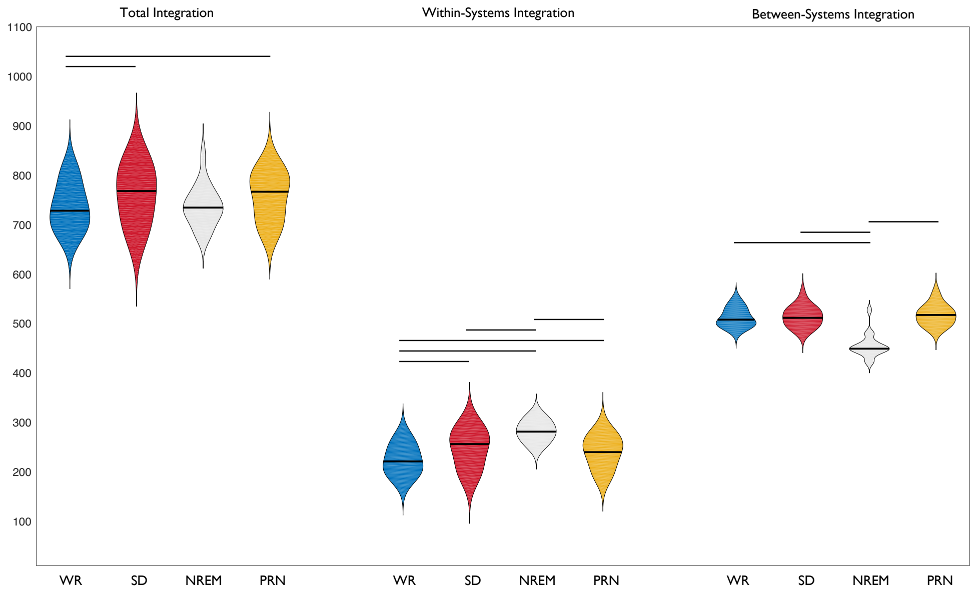


Fig S4 - There was a significant increase in total integration from RW to SD. This increase was due to an increase in within-systems (networks) integration, with no change in integration between networks. Alternatively, while there was no change in total integration during the NREM nap, there was a significant increase in within-systems and decrease in between-systems integration (solid lines represent significant differences between states).

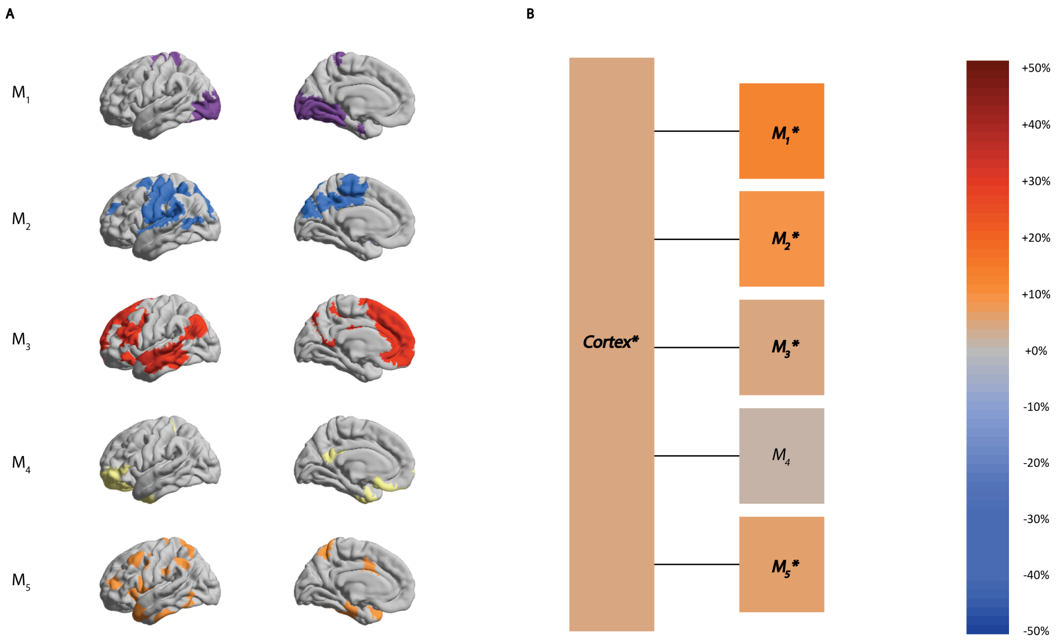


Fig. S5. **(A)** 5 networks (communities) were detected from the functional connectome in the WR condition using the Louvain modularity algorithm of the Brain Connectivity Toolbox. Visually, these appear to cover well described cortical networks in the literature: Visual; Somatomotor; Default Mode; Limbic; Attentional Networks. **(B)** When hierarchical integration was computed across these extracted networks, there was an observed increase from WR to the SD condition across the level of the entire cortex, and also within 4 of the 5 networks (*bold italics represent significant differences across states).

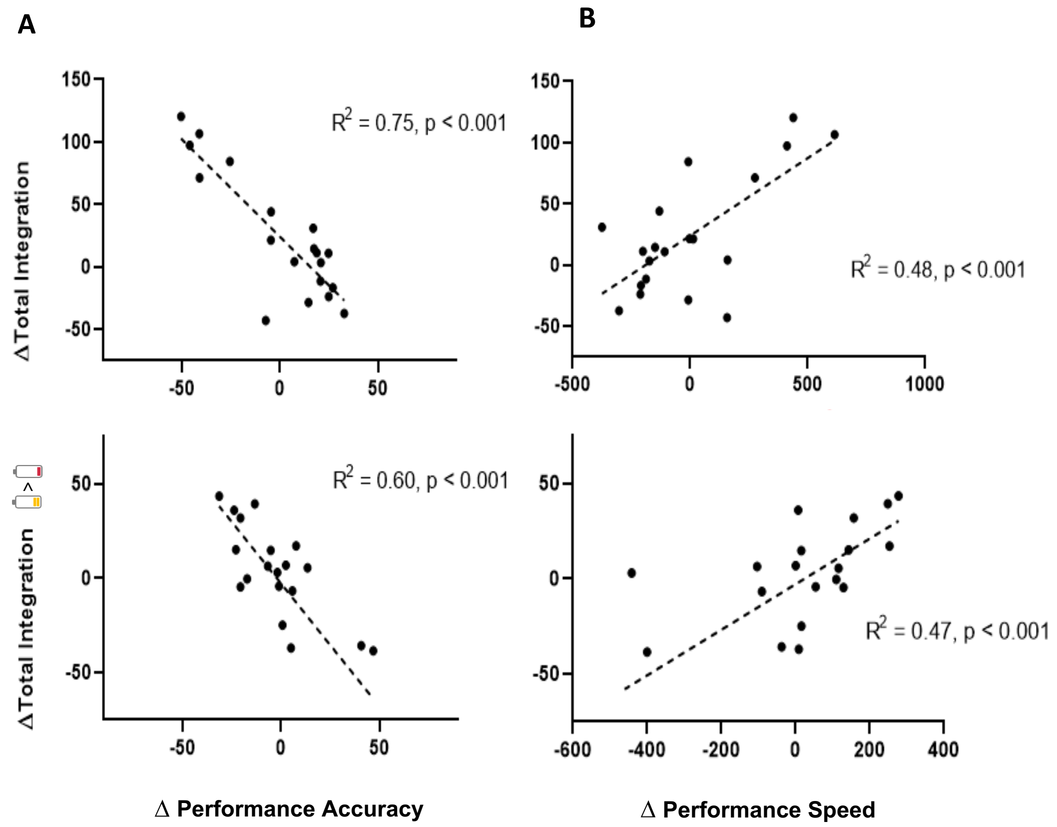


Fig. S6 – **(A)** Associations between the increase in total integration and cognitive performance. Similar to the functional clustering ratio, there was a significant negative relationship between change in integration and Accuracy and a positive relationship with Speed performance from RW to SD. **(B)** There was also a significant negative relationship between change in integration following the PRN and the change in performance from SD to PRN.

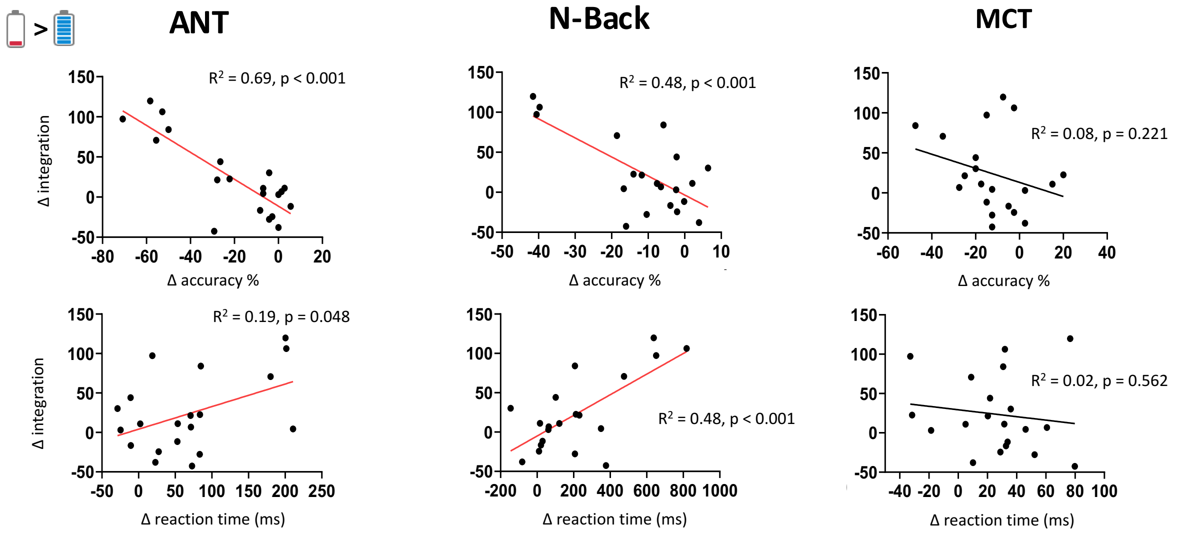


Fig. S7 - The change in performance from RW to SD on each task separately in comparison to the change in integration of cortical BOLD activity during each task. A greater increase in total integration was significantly related to worse Accuracy and Speed in the ANT and Nback tasks, but not the MCT.

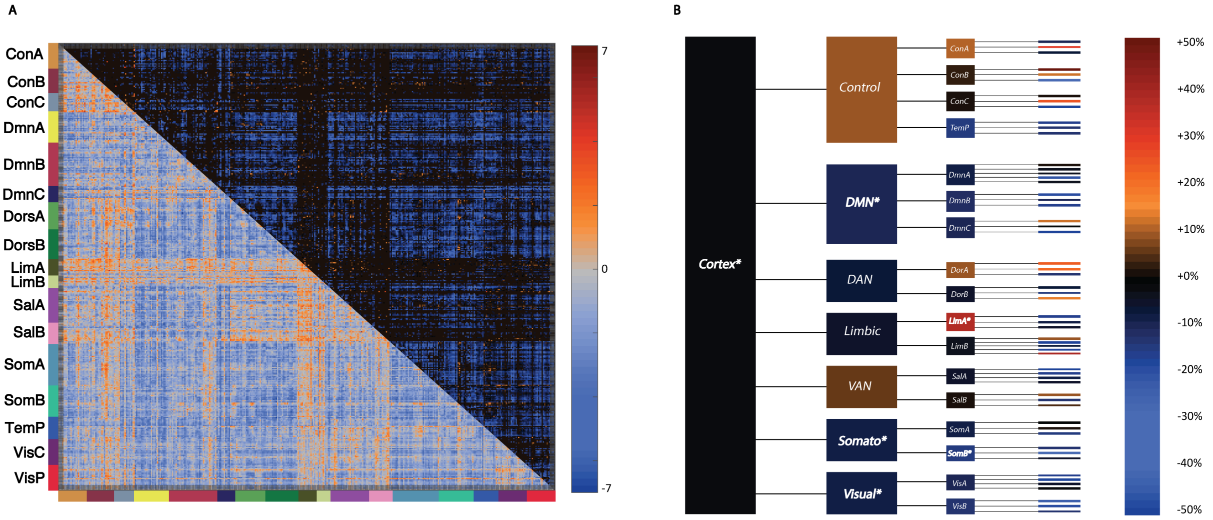


Fig. S8. **(A)** Overall a decrease in functional connectivity was observed from the SD to the PRN state. **(B)** Changes in integration are shown across different levels of a hierarchical model of the cortex: whole cortex, 7 networks, 17 networks, and 57 clusters. The total integration within the cortex decreased minimally from the WR to the SD state, and at the network level decreases were only observed within the Default Mode, Somatomotor and Visual networks (bold white italics depict significant changes, FDR corrected).
